# Taurine as a key intermediate for host-symbiont interaction in the tropical sponge *Ianthella basta*

**DOI:** 10.1101/2022.09.23.509140

**Authors:** Florian U. Moeller, Craig W. Herbold, Arno Schintlmeister, Maria Mooshammer, Cherie Motti, Faris Behnam, Margarete Watzka, Thomas Schweder, Mads Albertsen, Andreas Richter, Nicole S. Webster, Michael Wagner

## Abstract

Marine sponges are critical components of marine benthic fauna assemblages where their filter-feeding and reef-building capabilities provide bentho-pelagic coupling and crucial habitat. As potentially the oldest representation of a metazoan-microbe symbiosis, they also harbor dense, diverse, and species-specific communities of microbes, which are increasingly recognized for their contributions to dissolved organic matter (DOM) processing. Recent omics-based studies of marine sponge microbiomes have proposed numerous pathways of dissolved metabolite exchange between the host and symbionts within the context of the surrounding environment, but few studies have sought to experimentally interrogate these pathways. By using a combination of metaproteogenomics and laboratory incubations coupled with isotope-based functional assays, we showed that the dominant gammaproteobacterial symbiont ‘*Candidatus* Taurinisymbion ianthellae’ residing in the marine sponge, *Ianthella basta*, expresses a pathway for the import and dissimilation of taurine, a ubiquitously occurring sulfonate metabolite in marine sponges. ‘*Candidatus* Taurinisymbion ianthellae’ incorporates taurine-derived carbon and nitrogen while, at the same time, oxidizing the dissimilated sulfite into sulfate for export. Furthermore, we found that taurine-derived ammonia is exported by the symbiont for immediate oxidation by the dominant ammonia-oxidizing thaumarchaeal symbiont ‘*Candidatus* Nitrosospongia ianthellae’. Metaproteogenomic analyses also indicate that ‘*Candidatus* Taurinisymbion ianthellae’ likely imports DMSP and possesses both pathways for DMSP demethylation and cleavage, enabling it to use this compound as a carbon and sulfur source for biomass, as well as for energy conservation. These results highlight the important role of biogenic sulfur compounds in the interplay between *Ianthella basta* and its microbial symbionts.

## Introduction

Marine sponges are important components of coral reefs with a range of critical ecological functions [1]. Sponges play a crucial role in maintaining biomass in reef ecosystems by filtering dissolved organic matter (DOM) from large quantities of water and capturing it as biomass, which is then shed as particulate organic matter (POM) and passed to higher trophic levels [2–4]. Marine sponges often harbor dense, diverse and stable communities of microorganisms that are thought to play a role in DOM processing as well as a broad suite of other nutrient transformations [3, 5, 6]. These microbial symbionts have been shown to photosynthetically fix carbon [7], fix N2 [8], nitrify [9, 10], as well as carry out anammox, denitrification, and sulfate respiration [11, 12]. Furthermore, isotope labeling experiments have proven that symbiont-derived carbon and nitrogen can be found in sponge cells [7, 11, 13, 14], with more recent studies revealing an emerging view that not only microbial symbionts can incorporate DOM, but that sponge cells do as well, and can even translocate DOM-derived carbon and nitrogen to their symbionts [15–17].

Whilst the specificity, biogeography and environmental sensitivity of the relationship between sponges and their microbiome has been intensively studied [18–20], our understanding of the physiological interactions between sponges and their symbiotic microorganisms is still very limited. Genome-based investigations of the sponge microbiome have revealed an array of diverse metabolic potential encoded in symbiont genomes, including the processing of organic carbon, nitrogen, and sulfur compounds [21–24], autotrophic sulfur oxidation [25, 26], and vitamin synthesis [27, 28]. These metabolic processes may be mediated by a diverse array of transporters, encoded in the symbionts, to facilitate exchange with the host, as well as the surrounding environment (reviewed in [29]). In a few studies, metaproteomic analyses have revealed the expression of proteins indicative of metabolic interactions between the host and symbionts, including transport functions for typical sponge metabolites, and the potential mediation of host-symbiont interactions via the expression of eukaryotic-like proteins and cell-cell mediators [10, 30]. Despite these insights obtained from a variety of approaches, the high complexity of sponge holobionts, coupled with the inherent difficulty in culturing sponge symbionts, has made it challenging to irrevocably link host interactions and symbiont physiology with symbiont phylogeny.

One way of examining host-symbiont interaction, is by tracking compounds that may be translocated from the host to the symbiont, such as taurine. Taurine has been reported from a broad suite of sponge species, oftentimes as one of the most abundant free amino acids (up to 39% of the free amino acid pool; [31]). Free-living marine bacteria can utilize taurine produced by phytoplankton [32–34] and the genetic capacity for taurine utilization has been identified in many marine sponge microbiomes [25, 26, 35–37]. Taurine has been proposed to be an important intermediate driving metabolic exchange between marine bacteria and eukaryotic phytoplankton [32–34, 38], and yet, to date, there is no direct experimental evidence for the utilization of host-derived taurine by marine symbionts and of its role in mediating holobiont function, despite the abundance and widespread distribution of this compound in marine animals. Taurine, has however, been recently demonstrated to confer resistance to pathogens in mammalian gut systems, wherein taurine-metabolizing gut microbiota shield the host against infections by converting taurine to sulfide, an inhibitor of cellular respiration [39, 40]. Taurine is an ideal osmoregulator, as it is transported by a Na^+^-dependent transport system unique to β-amino acids and so, it is responsive to ionic activities and other osmotic agents [41, 42]. As a zwitterionic compound, extremely high intra-to extracellular concentration gradients for taurine can be maintained, and due to its lipophobic properties, it is not readily lost by diffusion [43]. Taurine also likely functions as a storage and transport compound, providing nitrogen and sulfur to symbionts in the gutless clams *Solemya reidi* [44] and *Solemya velum* [45]. The synthesis of sulfur-containing free amino acids has long been proposed as a strategy for sulfide detoxification and transport in chemoautotrophic symbioses [46–50] and taurine is the dominant free amino acid in many of these species [44, 46–50].

*Ianthella basta* [51] is a widely distributed sponge species found throughout the Indo-Pacific region [52]. In contrast to many marine sponge species with high microbial abundance of microbial symbionts [53], *I. basta* hosts only a low diversity of microbial symbionts and is populated by three dominant phylotypes including an Alphaproteobacterium, a Gammaproteobacterium and a Thaumarchaeote, which all occur at relatively high abundances in the sponge mesohyl [10, 54, 55]. This “low complexity” community is highly stable across different host morphotypes [56], across the large geographic range of the host, under different host health states [55] and across a wide suite of environmental conditions [57]. Thus, the *I. basta* holobiont represents a comparatively tractable model system for the investigation of sponge-symbiont physiological interactions. Recently, we were able to demonstrate that the thaumarchaeal symbiont of *I. basta* — ’*Candidatus* Nitrosospongia ianthellae’ — is an active ammonia oxidizer and that, surprisingly, this sponge does not harbor a bacterial nitrite oxidizer [10]. However, the physiology of the remaining two symbiont phylotypes remains unknown. In this study, we elucidate selected physiological traits of the gammaproteobacterial *I. basta* symbiont, which we named ’Candidatus Taurisymbion ianthellae’. Based on annotation of the metagenome-assembled genome (MAG) of this symbiont, we focused on the important biogenic sulfonate intermediate, taurine. We document the expression of a previously described [58], but infrequently observed pathway for the transport and dissimilation of taurine into sulfite, followed by sulfite oxidation into sulfate encoded by the symbiont, and provide experimental physiological support for this sponge-microbe interaction. For this purpose, we performed whole sponge incubations with added taurine and detected the predicted secreted end product, sulfate. We also employed the use of stable isotope-labeled taurine to track the incorporation of taurine-derived carbon and nitrogen into *I. basta* holobiont biomass, as well as the dissimilation of the nitrogen moiety to ammonium and its subsequent oxidation to nitrite.

## Materials and methods

### Sponge collection for metaproteogenomics and Ianthella basta holobiont taurine incubations

Two specimens of the purple thick morphotype of the sponge *Ianthella basta* were collected in waters offshore from Orpheus Island (18°36.878’S, 146°29.990’E), Queensland, Australia, in October 2010 and 2011, and processed as described in [10] for metaproteogenomic analysis. In addition, sponge explants were used for 48-hour batch incubations either in the presence or absence of taurine. The sponge explants used for the taurine incubations were derived from two large yellow thin morph *Ianthella basta* individuals (approx. 0.5 m x 0.5 m x 0.5 cm) collected from Davies Reef (18°49.354 S, 147°38.253 E) on 21^st^ November 2014 and 25^th^ August 2015. Upon collection, the individuals were maintained in flow-through reef seawater onboard the RV Cape Ferguson until its return to the Australian Institute of Marine Science (AIMS) where the individual was cut into small sponge explants between 10 cm x 10 cm x 0.5 cm and 5 cm x 5 cm x 0.5 cm and left to heal in AIMS aquaria overnight. Since *I. basta* individuals are thin, erect and laminar, while also exhibiting incurrent (ostia) and excurrent holes (oscula) that are spaced approximately 1 cm apart and on opposing sides of the sponge, relatively small samples can be representative of the entire aquiferous system [59]. The explants were prepared by cutting rectangular-like shapes, while preserving both sides of the fan-shaped sponge, thereby retaining multiple oscula and ostia on both sides. All explants were monitored at the beginning and end of the incubations for tissue health, and no major signs of tissue health regression or necrosis were observed. Furthermore, the edges of the cut sides are estimated to represent 4–8% of the volume of each explant (by assuming an edge thickness of 0.1 cm) and thus, the area of tissue wounds are small relative to the size of each explant. Live sponge explants were transferred to sulfate-free artificial seawater (SFASW; see Supplementary Note for media details) and rinsed 3 times to remove as much sulfate from seawater as possible. For all incubation experiments, *I. basta* explants were incubated at 25°C in the dark in continuously stirred acid-washed glass tanks either filled with 0.2 µm natural filtered seawater (FSW) or SFASW, and with or without taurine added at different time points and concentrations. All incubations were conducted in the dark to preclude light inhibition of nitrification. SFASW incubations were conducted to more readily discern the production of sulfate from the taurine-dissimilation/sulfite oxidation pathway, since natural seawater contains sulfate at high concentrations. In the first set of experiments, 1 mM unlabeled taurine was added at the beginning of the incubations and another 0.6 mM pulse was added at t = 36 h. For stable isotope incubations in standard seawater, ^15^N and ^13^C labeled taurine (2-Amino-^15^N-ethanesulfonic acid: Taurine-^15^N, 98 atom%; 2-Aminoethane-^13^C2-sulfonic Acid: Taurine-^13^C2, 99 atom%; Campro Scientific GmbH, Berlin, Germany) were added in equimolar concentrations (i.e. a total of 0.8 mM of ^15^N-taurine and 0.8 mM of ^13^C-taurine were added by the end of the incubations) simultaneously, so as to monitor potential incorporation into *I. basta* holobiont biomass as well as to track dissimilation into NH4, followed by successive oxidation to NO2 by the dominant thaumarchaeal symbiont [10]. For these two sets of experiments, incubations were conducted in closed 1 l acid-washed glass aquaria at room temperature and water sampling occurred at t = 0, 12, and 48 h, while sampling of *I. basta* biological material occurred at the end of the experiment at t = 48h. These two experiments utilized 12 explants (approx. 10 cm x 10 cm 0.5 cm; 9.14 ± 2.4 g sponge wet wt) from the individual collected in November 2014 (3 replicates per treatment with *I. basta* explants: SFASW + *I. basta*; SFASW + taurine + *I. basta*; FSW + *I. basta*; FSW + ^13^C^15^N-labeled taurine + *I. basta*). An additional follow-up experiment was conducted with explants derived from the individual collected in October 2015, in SFASW in closed 250 ml acid-washed glass aquaria but with higher temporal resolution (t = 0, 12, 24, 36, and 48 h) and 1 mM or 100 µM isotopically unlabelled taurine added at the beginning of the incubation. This set of experiments utilized 45 explants (approx. 5 cm x 5 cm 0.5 cm; 2.98 ± 0.82 g sponge wet wt), as each explant was sacrificially sampled at the end of each time point. For all sets of incubations parallel control incubations were performed under identical conditions, but without *I. basta* explants. All incubations were conducted in triplicates. We treated the explants from the same *I. basta* individual as non-technical replicates since differences in symbiont distribution within the relatively large sponge individuals used here, may occur.

For chemical analyses, seawater samples (10 ml) were taken from the containers (see above-described set of experiments) and filtered using 0.45 μm Sartorius Minisart cellulose acetate filters (Göttingen, Germany). Duplicate samples for dissolved inorganic nitrogen (NH4, NO2, NO3) were measured on a Seal AA3 segmented flow analyser and referenced against OSIL standards and in-house reference samples. Sulfate was measured via ion chromatography, followed by detection via suppressed conductivity. Confirmation of taurine uptake by *I. basta* was established following a two-step pre-column fluorometric derivatization HPLC method adapted from [60]. o-Phthalaldehyde-ethanethiol (OPA-ET; Sigma Aldrich; 750 μl), prepared daily in methanol (MeOH; Sharlau) buffered with 0.8 M borate (Sigma Aldrich; pH11) and aged at 4°C for 2 h, was added to 100 µl of seawater sample diluted in 900 µl MilliQ H2O. After 1 min, 50 μl of fluorenylmethyloxycarbonyl chloride (FMOC-Cl) [Sigma Aldrich], prepared in acetonitrile [Sharlau] and aged for at least 12 h, was added; the higher concentration of the derivatizing reagents was required to ensure complete reaction of the taurine (∼mM). After a further 1 min, 10 μl of the derivatized sample was injected onto an Agilent 1100 HPLC system (comprising a binary pump, autosampler, column oven and fluorescence detector, and operated using Chemstation for LC 3D Rev. A.10.01 and 735 Sampler Software v6.10). Samples (n = 2 samples per treatment per timepoint; n = 5 replicate injections per sample) were chromatographed on a 250 mm× 4.6 mm i.d., 5 μm particle size, HILIC (Phenomenex) column under the following conditions: 1 ml min^-1^ flow rate; column maintained at 40°C; mobile phases comprising MeOH-sodium acetate (0.36 M; pH 7.0)-H2O (55:8:37, v/v/v) (A) and acetonitrile-sodium acetate (0.36 M; pH 7.0)-H2O (10:5:85, v/v/v) (B); gradient 15% A at 0–2 min, 15–20% A at 2–6 min, 40% A at 6–9 min, 85% A at 9–17 min then held until 20 min; the column was flushed with 100% A at 20–21 min and held for a further 4 min before re-equilibration at 15% A for 5 min; fluorescence detection (excitation: 330 nm, emission: 460 nm). For calibration, a taurine stock solution (1 M; >99.9% Sigma Aldrich) was prepared in Milli-Q water and a dilution series prepared (10 mM–1 nM). γ-Aminobutyric acid (5 M GABA; Sigma Aldrich) was used as the internal standard. The dilution series was stored at 4°C. The calibration curve (R^2^ = 0.99) was used to quantify taurine in seawater. ^15^N-analyses of ammonium and nitrite from the incubation experiments with stable-isotope labeled taurine are described below.

### Metagenome assembly and genome binning

Sequencing libraries were prepared from the same DNA extracted from the individual collected in October 2010 and previously reported in [10] using the Nextera kit (Illumina) according to the manufacturer’s instructions and concentrations measured using the QuantIT kit (Molecular Probes, Life Technologies, Naerum, Denmark). The libraries were paired-end (2 × 150 bp) sequenced on an Illumina HiSeq2000 using the TruSeq PE Cluster Kit v3-cBot-HS and TruSeq SBS kit v.3-HS sequencing kit and on an Illumina MiSeq using v3 2 × 300 bp kits. Metagenome reads in fastq format, obtained from the Illumina sequencing runs, were end-trimmed at a minimum phred score of 15, a minimum length of 50 bp, allowing no ambiguous nucleotides and Illumina sequencing adaptors removed. Trimmed reads from each dataset were assembled using Spades version 3.11.0 [61], using default parameters and genomes were binned using Metabat v 2.12.0 [62]. MAGs from multiple Illumina datasets were dereplicated using dRep [63] and provisionally classified using CheckM [64]. A single high-quality gammaproteobacterial MAG was recovered after dereplication and uploaded to MaGe [65] for annotation. The mapped read pairs (at >98% identity) for the assembly of this MAG represented 3.02% of total QC paired reads and read coverage was approx. 26X. The MAG was assembled from run ERR884062 under BioProject PRJEB9378.

Two additional MAGs were assembled and binned from paired end reads from SRA runs derived from two sponge species, belonging to the same family as *Ianthella basta* (*Hexadella dedritifera* and *Hexadella* cf. *dedritifera*) [66]. Paired end reads for SRA runs SRR8088660, SRR8088661, SRR8088662 and SRR8088663 were downloaded from NCBI. BBduk (bbmap v. 37.61) was used to remove adapters and residual phiX (adapters.fa, phix174_ill.ref.fa.gz, ktrim=r k=21 mink=11 hdist=2), then quality filtered (minlen=99 qtrim=r trimq=15). Read sets were denoised using BayesHammer (SPAdes v3.15.3) [61] and merged using bbmerge (bbmap v. 37.61). Reads were assembled using SPAdes v3.15.3 (-k 21,33,55,77 --only-assemble) and metaspades (-k 21,33,55,77 --only-assemble) resulting in 8 assemblies (2 assemblers, 4 datasets). Each read set was mapped against each assembly using bbmap v. 37.61(minid=0.99 idfilter=0.99 ambiguous=best pairedonly=t killbadpairs=t mappedonly=t) for and MAGs were generated using metabat1 and metabat2 algorithms (metabat v 2.15). MAGs from each SRA run were then dereplicated with dRep v 1.4.3 dereplicate_wf (-comp 40 -con 10 -l 200000) [63]. “Winning” MAGs from each SRA run were combined and dereplicated using dRep v 1.4.3 (-comp 40 -con 10 -l 200000 -pa 0.6 --S_algorithm gANI --S_ani 0.965). Reads from each SRA run were remapped to each MAG using bbmap v. 37.61(minid=0.99 idfilter=0.99 ambiguous=best pairedonly=t killbadpairs=t mappedonly=t and relative abundance was calculated by dividing mapped fragments by total fragments for each metagenomic dataset.

### Cryosectioning and FISH for relative symbiont quantification

The aforementioned *I. basta* individual collected for metagenomic sequencing in October 2010 was assayed for its microbiological composition using fluorescence *in situ* hybridization (FISH). Briefly, after sample collection, the *I. basta* specimen was cut into tissue strips of approximately 4 cm^3^, fixed in 4% paraformaldehyde (PFA) for 1 h at room temperature and stored in ethyl alcohol (EtOH)-phosphate-buffered saline (PBS) (1:1) at -20°C. For FISH, PFA-fixed samples of *I. basta* were embedded in Neg-50 (Richard-Allan Scientific), a cryogenic water-soluble sectioning medium, and cut to 5-μm thin sections (Leica CM3050 S). We designed specific 16S rRNA-targeted FISH probes for the visualization of ‘*Ca*. Taurinisymbion ianthellae’ (GamD1137, 5’-CTC AAA GTC CCC GCC ATT-3’) and the dominant alphaproteobacterial symbiont (AlfD729, 5’-CGGACCTGGCGGCCGCTT-3’). To calculate the relative abundance of ‘*Ca*. Taurinisymbion ianthellae’, we applied the double-labeled FISH probe GamD1137 (Cy3), along with the double-labeled FISH probe for the α-symbiont (AlfD729 in Fluos) and the double-labeled probe mix EUB338-I, EUB338-II, and EUB338-III (all in Cy5; [67]) in equimolar concentrations. *I. basta* sponge section hybridizations were carried out in the presence of 25% formamide in the hybridization buffer (after formamide concentration optimization), and the stringency of the washing buffer was adjusted accordingly [68]. As a negative control, the nonsense probe nonEUB338-I (reverse complementary probe to EUB338-I; double labeled in FLUOS, Cy3, and Cy5) was applied on one section per well per slide hybridized [69]. Probe-conferred visualizations of fluorescent cells were carried out by a confocal laser scanning microscope (CLSM) (LSM 510 Meta; Zeiss, Oberkochen, Germany). Specifically, labelled bacterial cells were counted by eye in 10 randomly selected fields of view (each field contained an average of 260.6 ± 73 SD cells of ‘*Ca*. Taurinisymbion ianthellae’ at 630X magnification with an optical section thickness of 1 µm) on a 146.25 × 146.25 µm ocular grid. Based on these parameters and measurements we were able to estimate the average density of ‘*Ca*. Taurinisymbion ianthellae’ within the *I. basta* mesohyl as well as the fraction of ‘*Ca*. Taurinisymbion ianthellae’ relative to total bacterial cells stained by the EUB338 probe mix.

### DNA extraction from whole sponge tissue and qPCR for symbiont quantification

Between 80–150 mg of *I. basta* sponge tissue was thawed (if previously frozen at -80 °C), rinsed successively (3x) in 1X calcium- and magnesium-free artificial seawater (CMF-ASW; see Supplementary Note for media details) and immediately ground into a paste with a mortar and pestle in liquid N2. After resuspension in TE buffer, DNA was extracted from the suspension using the same methods as described in [10] for DNA extraction prior to symbiont quantification via quantitative PCR (qPCR). DNA was extracted from the two individuals used for metaproteogenomics (see above) as well as three additional yellow thin morphotype healthy individuals from a previous study examining the persistence of microbial symbionts in diseased and healthy specimens of *I. basta* (Luter *et al*., 2010). qPCR was used to quantify the number of 16S rRNA genes of ‘*Ca*. Taurinisymbion ianthellae’, the thaumarchaeal ammonia-oxidizer ‘*Ca.* Nitrosospongia ianthellae’, and the dominant α-proteobacterial symbiont using specific primers designed for each symbiont phylotype in [10].

### Phylogenetic analyses

Sequences for the ‘*Ca*. Taurinisymbion ianthellae’ 16S rRNA species tree were assembled by merging a subset of sequences described by [70], along with top BLAST hits from representative gammaproteobacterial sequences, and the 16S rRNA sequences from the most closely related members with sequenced genomes. These sequences were aligned via the SINA aligner and the phylogenetic trees constructed in IQ-Tree, version 1.6.2 [71] using the model TN+F+R6 (after model selection using Model Finder; [72] with 10,000 ultrafast bootstraps (UFBoot, [73]). In addition, 16S rRNA gene sequences related to ‘*Ca*. Taurinisymbion ianthellae’ were identified in short read archive (SRA) datasets using IMNGS (www.imngs.org, [74]) with default parameters. Successfully mapped reads were then aligned to the reference alignment using SINA and placed into the reference tree using the Evolutionary Placement Algorithm (EPA; [75]) implemented in RAxML-HPC 8.2.11 [76]. Phylogenomic reconstruction was based on a concatenated amino-acid alignment of 43 marker genes constructed with CheckM [64] and trees were constructed using the model LG+F+R7 (after model selection using Model Finder; [72]) with 1000 ultrafast bootstraps (UFBoot, [73]).

### Proteomics

Proteome analysis was performed as done previously [10]. In brief, protein extracts were separated by 1D PAGE followed by liquid chromatography-based mass spectrometry (1D-PAGE-LC-MS/MS). MS spectra and MS/MS spectra were acquired from eluting peptides in a LTQ Orbitrap Velos hybrid mass spectrometer (Thermo Fisher Scientific, Waltham, MA, USA), as described previously [77, 78] and searched against predicted protein sequence databases composed of the *I. basta* symbiont-enriched metagenome bins and common laboratory contaminants using the Sorcerer SEQUEST (v.27, rev. 11) algorithm. Protein identifications were filtered with Scaffold version 3.5.1 as described previously [79]. For protein identification only peptides identified with high mass accuracy (maximum ± 10 ppm difference between calculated and observed mass) were considered and at least two exclusively unique peptides were required to identify a protein. False-discovery rates (FDRs) were estimated with searches against a target-decoy database [80, 81]. For relative quantitation of proteins, normalized spectral abundance factors (NSAF) were calculated for each sample according to the method of [82] and averaged for all replicates and samples. The proteomics raw data have been deposited to the ProteomeXchange Consortium via the PRIDE partner repository with the dataset identifier PXD012484 and 10.6019/PXD012484.

### NMR-, LC-MS-based detection and quantification of taurine in *Ianthella basta*

To establish the presence of taurine in *I. basta*, a section of tissue from a yellow thin morphotype was excised by scalpel, weighed wet, and exhaustively extracted with 90% MeOH in MilliQ water. The MeOH extract was lyophilized overnight (Dynavac Freeze-drier). The dry MeOH extract (∼5 mg) was dissolved in 1 ml deuterated methanol (CD3OD; Cambridge Isotope Laboratories) for analysis by proton nuclear magnetic resonance (^1^H NMR; Bruker Avance 600 MHz, 5 mm CPTXI inverse ^1^H-^13^C/^15^N Z-gradient cryogenically cooled probe, referenced to CD3OD 3.31 ppm, standard 1D zg pulse sequence). Given the crude nature of the extract and the likelihood that the diagnostic signals would be unable to be distinguished from similar chemistry in the extract, 20 mg (dissolved in ∼1 ml 50% MeOH) was chromatographed over a pre-equilibrated Strata C18-E 55 μm, 70Å, 1000 mg solid phase extraction cartridge (C18-SPE; Phenomenex) and eluted with 10 ml of H2O, 10% MeOH, 50% MeOH and 100% MeOH. The four fractions (Fractions 1–4) were evaporated under nitrogen stream at 40°C and lyophilized to dryness. Fractions 1–2 were prepared in 100% deuterated H2O (D2O; Cambridge Isotope Laboratories; referenced at 4.39 ppm) and Fractions 3–4 in 10% CD3OD: 90% CD3OD (referenced to 3.31 ppm) and analysed by ^1^H NMR. ^1^H and COSY NMR spectra of 100 μM taurine standard, prepared in D2O, was used to confirm identification of taurine in Fraction 1. Final confirmation was provided by spiking the ^1^H NMR sample with 32 μl and 128 μl 100 μM taurine and measuring the increase via integration of the two triplet signals.

To corroborate the NMR assignment, Fraction 1 was also analysed by LC-MS. A 10 μl aliquot of the aqueous Fraction 1 was injected onto an Agilent 1100 HPLC system coupled to a Bruker Esquire 3000 Quadrupole Ion Trap LC-MS and eluted from a 150 mm x 3 mm Luna 5μ NH2 (Phenomenex) column under the following conditions: 0.8 ml min^-1^ flow rate; column maintained at 40°C; mobile phases comprised of H2O + 0.1 % formic acid (A) and MeOH + 0.1% formic acid (B); isocratic elution at 90% A: 10% B for 10 min. The ionspray voltage was set to -4500 V, declustering potential -500V, curtain gas flow 9 l min^-1^, ion source gas 40 PSI, and spray temperature at 350°C. Data was acquired over the mass range 70–200 *m/z*; product ion MS/MS data was acquired over 50–200 *m/z* to establish the fragmentation pattern. Retention time and total and extracted ion chromatographs (positive mode) were compared against a 100 μM taurine standard prepared in MilliQ.

The concentration of taurine in sponge tissue was measured by quantitative NMR (qNMR) following [83]. Briefly, five sponge explant samples were extracted and chromatographed over C18–SPE as described above. ^1^H NMR spectra of Fraction 1 (∼16 mg dry extract) from each of the five sponge extracts were acquired in 800 μl D2O (δH 4.39) at 298 K, with sweep width 12 ppm (7184 Hz), 90° pulse, 35 sec relaxation delay, receiver gain 16, 2 dummy scans, 16 acquisition scans and 64 k data points. The external reference signal was calibrated using a stock solution of 2 mM taurine in D2O. The concentration of taurine in the NMR samples was determined by comparing the signal integrals of well resolved non-exchangeable protons (2H) centred at 3.26 ppm in a 0.20 ppm window with that of the reference signal. Concentration was normalized to wet weight of sponge (wet wt).

### Isotopic analysis of whole sponge tissue, total nucleic acids, and ^15^N in NH4 & NO2 using EA-IRMS

For the determination of ^13^C- and ^15^N-enrichment within the holobiont, sponge tissue samples were rinsed briefly 3X in 1X CMF-ASW, freeze-dried, ground to a fine powder in a mortar, and stored dry prior to analysis. δ^13^C- and δ^15^N-values of sponge tissues were determined by an elemental analyzer (EA 1110, CE Instruments, Milan, Italy) coupled to an isotope ratio mass spectrometer (IRMS; Finnigan MAT Delta^Plus^ IRMS with a Finnigan MAT ConFlo III interface). For determination of ^13^C- and ^15^N-enrichment in total nucleic acids, *I. basta* whole sponge tissue was homogenized using the above procedure employing liquid N2, followed by nucleic acid extraction according to the procedures outlined in [84]. The extracted nucleic acids were then dried overnight in an oven, and δ^13^C- and δ^15^N-values were determined via the same procedure as above. For quantification of the amount of ^13^C- and ^15^N-derived taurine that was incorporated into *I. basta* holobiont biomass, the following formulas were used:

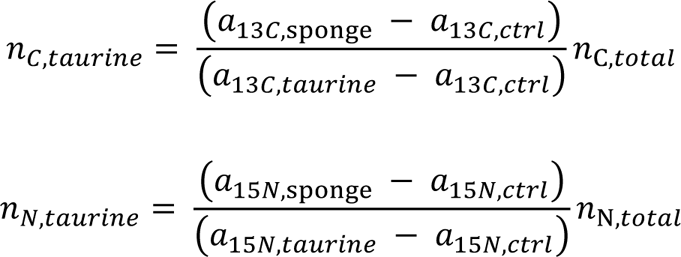

where the displayed symbols, abbreviations and designations refer to: *a* — isotope fraction ^13^C/(^12^C+^13^C) or ^15^N/(^14^N+^15^N) given in at%; *n*: number of carbon or nitrogen elemental atoms; *sponge* — sponge material; *ctrl* — natural isotope abundance control; *total* — total amount of carbon or nitrogen in the analyzed sample. The denominator in both equations (*a*15_*C,N,taurine*_ − *a*15_*C,N,Ctri*_) were calculated via a standard two-pool mixing model [85]. For comparability of the measurement values obtained from distinct incubations, *n* was normalized to the wet weight of the utilized *I. basta* explants.

For ^15^N-analysis of NH4 and NO2, sampled seawater from the glass tanks with the labeled taurine incubations was filtered through pre-combusted GF/Fs (Whatman International; treated for 4 h at 450°C), and subsequently through 0.2 µm polycarbonate filters (Sartorius). All samples were immediately frozen at -20 °C for later analysis. Prior to shipment to the University of Vienna, samples were thawed at room temperature and the microbial inhibitor phenylmercuric acetate was added (to a final concentration of 10 µM). Upon arrival in Vienna the samples were promptly stored at -80°C. For determination of the ^15^N content in ammonia, NH4 was extracted from samples via microdiffusion as described in [10] and following the protocol from [86]. Briefly, 100 mg MgO and an acid trap (acidified cellulose filter disc enclosed in a semi-permeable Teflon membrane) were added to 9 ml of sample and 1.5 ml of 3 M KCl. After 5 days shaking at 35°C, the acid traps were removed, dried over concentrated sulfuric acid and analyzed for the ^15^N content by an elemental analyzer (EA 1110, CE Instruments, Milan, Italy) coupled to an isotope ratio mass spectrometer (IRMS; Finnigan MAT Delta^Plus^ IRMS with a Finnigan MAT ConFlo III interface). The nitrogen isotope composition of NO2 was determined by a method based on the reduction of NO2 to N2O by using sodium azide under acidified conditions following the protocol of [87]. Briefly, 1 ml sample or standard was transferred to 12 ml exetainer and 1ml 1M HCl was added. After purging the vials with helium to expel air-N2O from the sample headspace, 150 µl 1M sodium azide buffer (in 10% acetic acid solution) was injected and the vials were placed on a shaker at 37 °C for 18 h. The reaction was quenched by injecting 250 µl of 10M NaOH. Derived N2O was analyzed using a purge-and-trap GC/IRMS system (PreCon, GasBench II headspace analyzer, Delta Advantage V IRMS; Thermo Fischer). Equivalently to the above calculations, the amount of taurine-derived NH4 and NO2 were calculated as follows:

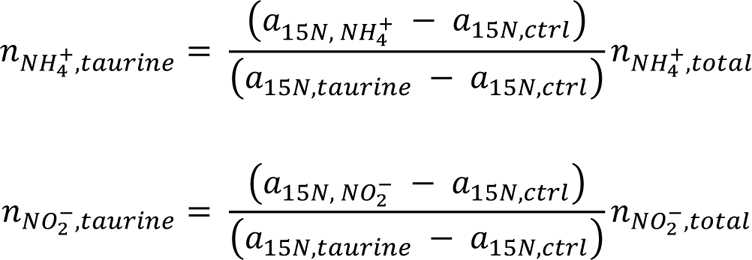

By taking into account that the isotopically labeled nitrite emerges from oxidation of ammonia obtained from taurine dissimilation, the total amount of NH4 released by the holobiont through taurine metabolization was calculated via:

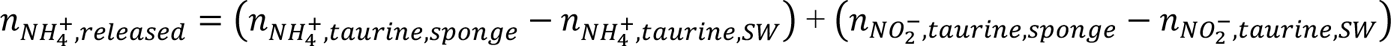

where SW refers to the values determined from control incubations utilizing seawater without sponge explants. For comparability of the measurement values obtained from distinct incubations, *n* was normalized to the wet weight of the utilized *I. basta* explants.

### NanoSIMS analysis of Ianthella basta

NanoSIMS measurements were undertaken on two of the three *I. basta* explant samples obtained from the ^13^C^15^N-taurine incubations. Specimens were selected for maximum isotope enrichment, based on results from EA-IRMS bulk analysis (0.62 and 0.55 at% ^15^N; 1.19 and 1.15 at% ^13^C). Briefly, approximately 0.5 cm^3^ sections were cut from *I. basta* samples, which had been fixed in 4% PFA and stored in a 1:1 mixture of PBS and 96% ethanol at -20° C. The thus obtained specimens were then rinsed successively (3x) in 1X calcium- and magnesium-free artificial seawater (CMF-ASW) and gently dissociated in 1X CMF-ASW with a scalpel, mortar and pestle, and a final step of sonication for two 30 second cycles at 35% power using a sonication probe [Sonopuls HD2200 equipped with probe MS73 (output frequency 20 kHz, power 200 W, amplitude: 310 μm), Bandelin, Germany]. The suspensions were then filtered through 0.45 μm cellulose acetate filters (type Minisart, Sartorius, Göttingen, Germany) to remove a large portion of remaining sponge cells and nuclei (microscopic visualizations showed a lack of sponge nuclei after filtration), centrifuged at 10,844 × g for 10 min at 4°C to pellet cells, and resuspended in 50% EtOH/50% 1X PBS solution. To visualize all DNA-containing cells present in the sample, a 30 µL aliquot of the suspension was spotted onto antimony-doped silicon wafer platelets (7.1 x 7.1 x 0.75 mm; Active Business Company, Germany) and dried at 46° C, before staining with a 0.1 % (w/v) solution of the DNA-binding dye 4,6-diamidino-2-phenylindole (DAPI) (Sigma, Deisenhofen, Germany) at room temperature for 5 min. Excess DAPI was removed by rinsing with distilled water and the slides were air-dried. Samples were then imaged on an epi-fluorescence laser microdissection microscope (LMD, Leica LMD 7000) using a 40× air objective for pre-selection of suitable NanoSIMS measurement areas, which were subsequently marked utilizing the LMD laser. High resolution imaging of fluorescent cells in marked regions was carried out by a confocal laser scanning microscope (CLSM) (LSM 510 Meta; Zeiss, Oberkochen, Germany). NanoSIMS measurements were performed on a NanoSIMS 50L (Cameca, Gennevilliers, France) at the Large-Instrument Facility for Environmental and Isotope Mass Spectrometry at the University of Vienna. Further details on NanoSIMS data acquisition and analyses are presented in the Supplementary Information.

## Results and Discussion

### *A metagenome assembled genome and phylogenetic analyses of the dominant γ-proteobacterial symbiont of* Ianthella basta

From the extensive metagenomic data set used in [10] for genomically characterizing the thaumarchaeal symbiont of *I. basta*, we retrieved a 2.35 Mbp MAG consisting of 97 contigs, representing a nearly complete genome (94.8%) with a very low level of contamination (Table S1). Phylogenetic analysis of the only 16S rRNA gene copy in the MAG demonstrated that this gene specifically clustered at 99% nucleotide identity with previously recovered sequences of the dominant gammaproteobacterial symbionts from *I. basta* (Fig. S1) [10, 55, 56] and fall into the neighborhood of sponge specific clusters 144–148 (SC) [70]. A phylogenomic reconstruction using concatenated single copy conserved marker genes (Fig. 1) [64], confirmed the affiliation of this MAG with the Gammaproteobacteria and revealed that it can be further classified into the Arenicellales order and within the LS-SOB family (originally named after the bacterial MAG extracted from the glass sponge *Lophophysema eversa,* which encodes a sulfur oxidation pathway [88]; based on the Genome Taxonomy Database [89]).

**Figure 1.**
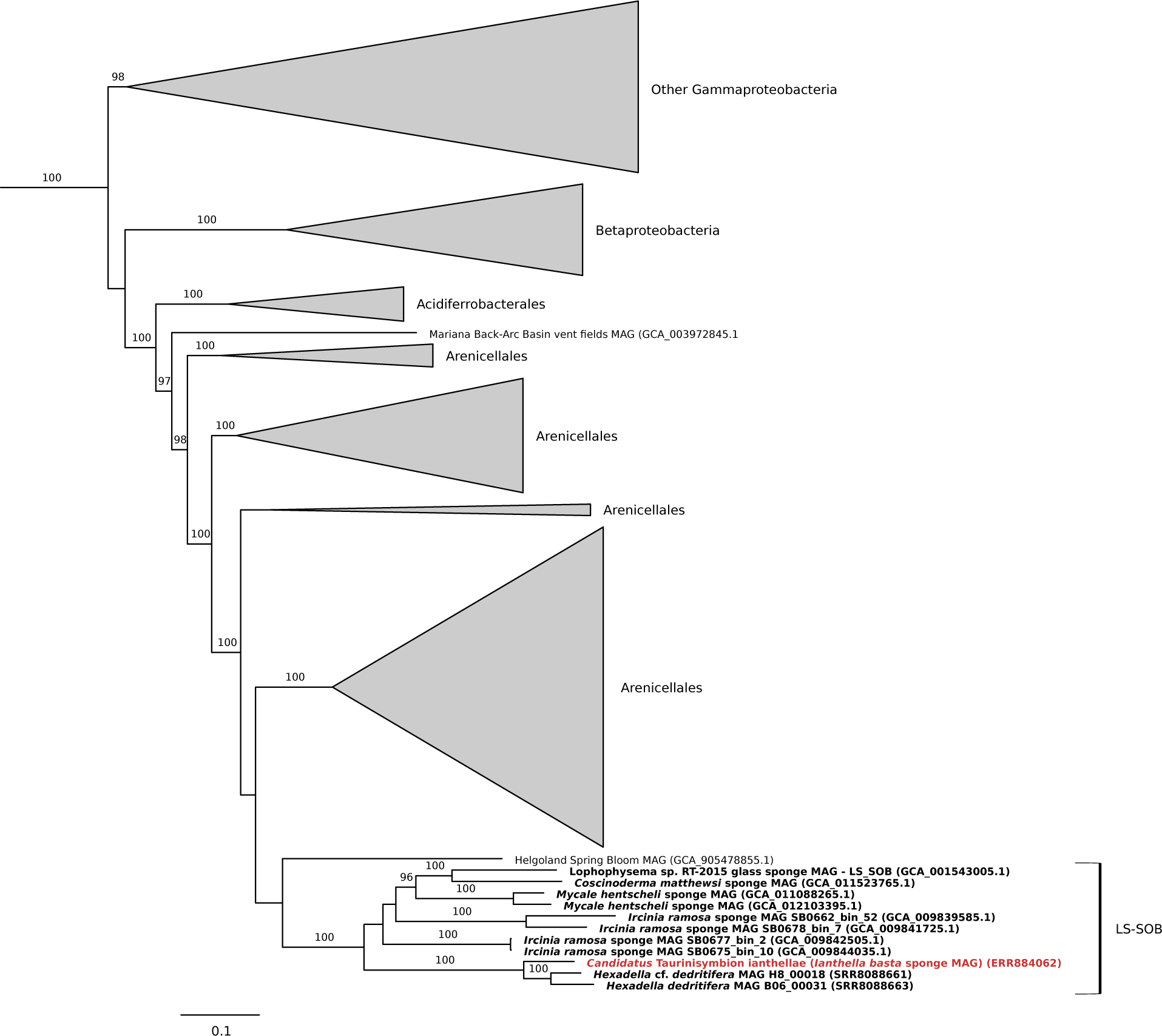
Maximum-likelihood phylogenomic tree (after automatic model selection with IQ-Tree) of ‘*Ca*. Taurinisymbion ianthellae’ (red and bold) along with other selected proteobacterial symbionts (sponge symbionts in bold font). The phylogenomic tree is based on 43 concatenated universal, single-copy marker genes identified with CheckM.

Based on the concatenated gene tree analyses, we further queried representative 16S rRNA gene sequences pulled from LS-SOB genome bins, against the IMNGS 16S rRNA gene amplicon database [74], and placed the hits into the reference 16S rRNA gene tree using the Evolutionary Placement Algorithm [75]. As a result, 3 sets of short reads were placed into the branch leading to the dominant γ-proteobacterial symbiont sequences obtained from *I. basta* (Fig. S1). The closest two groups (I4, I6) comprised 5 (out of 6 total) OTU reads derived from unspecified coral and sponge samples, 4 of which were OTU reads from corals sampled from the same study site as *I. basta* for this study (Orpheus Island, Queensland, Australia). 141 OTU reads were placed into the I7 group, of which 29 and 4 were derived from sponge and coral metagenomes, respectively (the other 112 OTU reads were derived from other metagenomic data sets spanning a variety of environmental matrices). Most interestingly, out of this group, three OTU reads were obtained from two sympatric marine sponge species (NE Atlantic) within the same genus, (*Hexadella dedritifera* and *Hexadella* cf. *dedritifera*), both of which are also affiliated with the same sponge family as *I. basta*, the Ianthellidae. These two closely related phylotypes, from *H. dedritifera* and *H*. cf. *detritifera*, respectively, exhibited 16S rRNA gene identities of ∼97% (over 418 bp) with each other, 95–97% with the *I. basta* γ-symbiont, and were found to comprise 33–42% of the total amplicon dataset derived from these sponges [90]. Noting that these two Hexadella species harbor abundant and closely related γ-phylotypes, and also contain AOA-symbionts closely related to the AOA in *I. basta* [66], we assembled and binned two new γ-symbiont MAGs from the published raw read data set of these two Hexadella sponges.

The phylogenomic reconstruction confirmed that these two Hexadella γ-symbiont MAGs were most closely related to the *I. basta* γ-symbiont (Fig. 1), and AAI analyses indicated that the *I. basta* γ-symbiont shares AAI values of 76–80% with these two Hexadella γ-symbionts. Among the other LS-SOB genomes retrieved from marine sponges [37, 88, 91–93], the *I. basta* γ-symbiont shares AAI values of 54–59%. Since these identity values signify that the γ-symbiont represents a new species within a new genus along with the other two Hexadella γ-symbionts — 60–80% AAI is typical for organisms grouped at the genus level [94, 95] — we propose the name ‘*Candidatus* Taurinisymbion ianthellae’. Taurinisymbion gen. nov. (Tau.ri.ni.sym’bi.on. N.L. n. taurinum, taurin; Gr. pref. sym, together; Gr. masc. part. n. bion, a living being; N.L. masc. n. Taurinisymbion, a taurin-degrading living partner) for the *I. basta* γ-symbiont. The name *Taurinisymbion* describes this organism’s ability to utilize taurine as a source of energy and biomass whilst residing in symbiosis within a sponge, and we were able to identify the same pathways encoded by ‘*Ca*. Taurinisymbion ianthellae’ for taurine utilization in the two Hexadella γ-symbiont MAGs, thereby suggesting a conserved evolutionary function that may be adaptive for this proposed genus (see below; Table S2). The species name *ianthellae* refers to its discovery and description as a symbiont of the marine sponge, *I. basta*.

### ’Candidatus Taurisymbion ianthellae’ is an abundant symbiont of *Ianthella basta*

Consistent with the high coverage of reads and peptides assigned to ‘*Ca*. Taurinisymbion ianthellae’ in the metaprotoegenomic data sets from *I. basta* and the dominance of this phylotype in other *I. basta* studies [54–56], we observed a high abundance of this symbiont (Fig. 2) in the mesohyl of *I. basta* using fluorescence *in situ* hybridization (FISH) with a newly designed specific 16S rRNA-targeted probe. Quantitative FISH revealed that ‘*Ca*. Taurinisymbion ianthellae’ achieved average cell densities of 1.22 ± 0.34 (SD) × 10^10^ cells cm^-3^ sponge and accounted for 24 ± 5.6% (SD) of all cells hybridizing with the probe set EUB338-I-III targeting most bacteria (which does not cover the thaumarchaeal symbiont of *I. basta*). Moreover, absolute quantification of ‘*Ca*. Taurinisymbion ianthellae’ using qPCR with newly designed symbiont specific primers, showed an average absolute abundance of 2.78 ± 1.5 (SD) × 10^10^ 16S rRNA gene copies g^-1^ sponge wet wt, which is in the same range as the population density values observed from the FISH images (assuming that 1.2 g sponge wet wt = 1 cm^-3^; [96]). The absolute abundances obtained for the dominant thaumarchaeal and alphaproteobacterial symbionts are also in the 10^10^ 16S rRNA gene copies g^-1^ sponge wet wt range (Fig. S2). The ratio of ‘*Ca*. Taurinisymbion ianthellae’ 16S rRNA sequences to the total 16S rRNA genes derived from qPCR assays (targeting the γ-, α-, and thaumarchaeal symbionts of *I. basta*) was 41.1 ± 34 (SD)%. Nevertheless, it should be noted that qPCR abundances cannot directly be translated to cell numbers as (i) symbionts can have multiple rRNA gene copies (although the ‘*Ca*. Taurinisymbion ianthellae’ MAG has only one 16S rRNA gene copy), (ii) genome polyploidy might occur in sponge symbionts and (iii) extracellular DNA of the symbionts will also be detected. Similarly, using FISH for quantification may undercount microbial cells with low ribosome contents.

**Figure 2.**
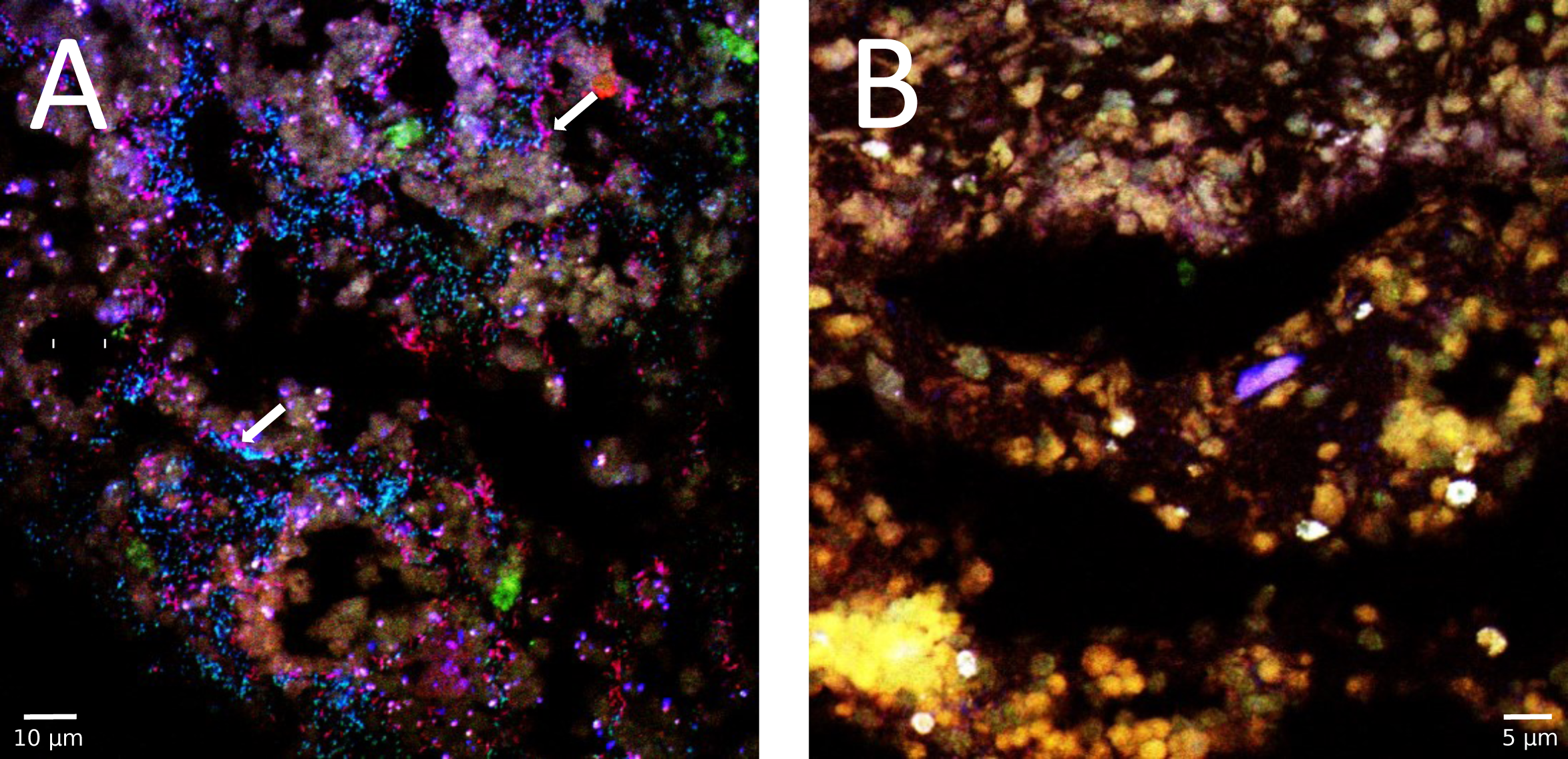
Fluorescence *in situ* hybridization of 5 μm thick cryo-sections of *Ianthella basta* using double-labeled probe sets [55]. A) ‘*Ca.* Taurinisymbion ianthellae’ (red), *Alphaproteobacteria* symbiont*-*probe (green), and *Bacteria-*probe (EUB338-I-III; blue); B) negative controls using the double-labeled non-EUB338 probe set are depicted in all three color channels. The combination of the ‘*Ca.* Taurinisymbion ianthellae’ specific probe signals in red with the general *Bacteria* probe signals in blue, results in the visualization of ‘*Ca.* Taurinisymbion ianthellae’ as magenta signals and representative cells are indicated with white arrows. The α-symbiont appears in cyan. Green structures represent autofluorescence.

### ‘Ca. Taurinisymbion ianthellae’ *expresses proteins for taurine catabolism and subsequent NH4 and SO4 export*

To analyse the metabolic potential of ‘*Ca*. Taurinisymbion ianthellae’, we annotated its MAG by employing the metaproteogenomic data set presented in [10] and were able to assign 149 proteins to this symbiont MAG. These proteins represented 35.7%, in NSAF values, of all proteins assigned to the *I. basta* microbiome (Table S3), highlighting the physiological dominance of ‘*Ca*. Taurinisymbion ianthellae’. From this metaproteogenomic dataset, we concluded that ‘*Ca*. Taurinisymbion ianthellae’ is a facultative anaerobe with an ability to metabolize a variety of organic compounds. The genomic potential for the major central metabolic pathways, such as glycolysis, the pentose phosphate pathway, the tricarboxylic acid (TCA) cycle, the glyoxylate bypass, a conventional respiratory chain (complex I -IV; two respiratory complex IV were found to be encoded, a *cbb*3- and a *aa*3-type cytochrome c oxidase; complex V), as well as the genes for denitrification of NO2 to N2O, and DMSP catabolism, were all present in the recovered MAG (Fig. 3, Table S2). A further discussion of a few of these genomic features, with a focus on DMSP metabolism, is provided in the Supplementary Information.

**Figure 3.**
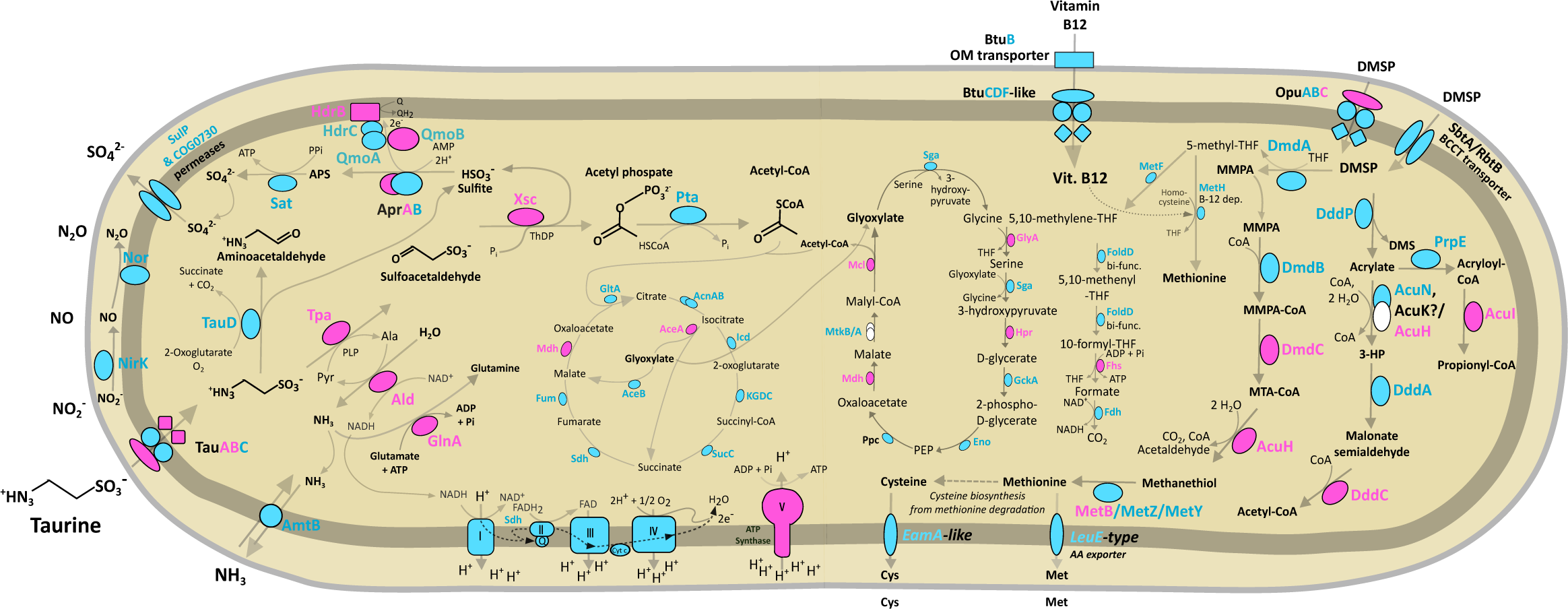
Taurine and DMSP metabolic pathways identified and expressed in ‘*Ca.* Taurinisymbion ianthellae’. Central carbon and nitrogen metabolism in ‘*Ca.* Taurinisymbion ianthellae’ based on metaproteogenomic analyses with a focus on taurine and DMSP metabolism pathways along with the relevant import and export of exchanged metabolites. Depicted are proteins necessary for taurine import, followed by taurine dissimilation, energy conservation via sulfite oxidation and the electron transport chain, along with carbon incorporation via the glyoxylate bypass. Similarly depicted are the proteins necessary for DMSP import as well as the corresponding DMSP cleavage and demethylation pathways, including the oxidation of 5-methyl-THF and its role as a carbon donor for the conversion of glycine to serine. Enzymes detected via metaproteomics are highlighted in pink, while enzymes only identified in the ‘*Ca.* Taurinisymbion ianthellae’ MAG are highlighted in turquoise. Missing pathway genes are depicted as white shapes. AMP, adenosine monophosphate; APS, adenosine-5’-phosphosulfate, Cys, cysteine; DMSP, dimethylsulfoniopropionate; DMS, dimethylsulfide; 3-HP, 3-hydroxypropanoate; Met, methionine; MMPA, methylmercaptopropionate; MTA-CoA, methylthioacryloyl-Coenzyme A; PEP, phosphoenolpyruvate; PLP, pyridoxal 5’-phosphate; Q, quinone; ThDP, thiamine diphosphate; THF, tetrahydrofolate. Additional information for genes and pathways depicted in the figure can be found in Supplementary Tables S2–S5.

Most notably, ‘*Ca*. Taurinisymbion ianthellae’ encodes a previously described pathway for taurine import and transamination (*tauABC, tpa, xsc*) [97] into sulfite via a desulfonated intermediate sulfoacetaldehyde, as well as a pathway for the subsequent oxidation of the resultant toxic and highly reactive sulfite into sulfate via the cytoplasmic enzymes APS reductase (a*prAB*) and ATP sulfurylase (*sat*) [58]. This pathway consumes AMP and sulfite, and releases ATP via substrate-level phosphorylation and sulfate, while channeling two electrons from the oxidation of sulfite, with its very negative redox potential, to the quinone pool via the membrane-bound QmoAB. Consistent with other sulfur oxidizing bacterial genomes, the *qmoC* gene is replaced by *hdrBC* genes ([98] and references therein), with the *hdrBC* genes found to be adjacent to *qmoAB* in ‘*Ca*. Taurinisymbion ianthellae’, and the HdrB subunit expressed in the holobiont (Fig. 3, Tables S2 & S3). For the potential export of sulfate, two putative sulfate permeases (COG0730 permease family, [99]; COG0659, SulP family permease), were detected in the ‘*Ca*. Taurinisymbion ianthellae’ MAG. Interestingly, the gene encoding taurine dioxygenase (*tauD*), an alpha-ketoglutarate-dependent dioxygenase, which desulfonates taurine into aminoacetaldehyde (the fate of which is currently unknown) and sulfite, was also found encoded by ‘*Ca*. Taurinisymbion ianthellae’. TauD is involved in assimilation of sulfur from taurine in several aerobic bacteria [100]. Furthermore, we found many of these genes (*tauABC*, *tpa*, *tauD*, *xsc*, *aprAB*, *qmoABC*) encoded in the previously mentioned Hexadella γ-symbiont MAGs, which belong to the same proposed genus as ‘*Ca*. Taurinisymbion ianthellae’ (Table S2).

Several of the genes involved in taurine degradation were also found to be expressed after being mapped to the ‘*Ca*. Taurinisymbion ianthellae’ MAG. Specifically, we detected the expression of the taurine transporter TauABC (with TauAB detected only when the 454 metagenome dataset was included as a database for initial metaproteomic analyses), as well as the two enzymes, taurine-pyruvate aminotransferase (Tpa) and sulfoacetaldehyde acetyltransferase (Xsc), involved in the sequential dissimilation of taurine to sulfite via sulfoacetaldehyde. Additionally, 3 out of 7 proteins necessary for the subsequent oxidation of sulfite to sulfate (AprA, QmoB and HdrB; while Sat was not detected in the metaproteome data set) were expressed. A phosphate acetyltransferase is present in the ‘*Ca*. Taurinisymbion ianthellae’ MAG, which would convert the acetyl phosphate produced from the activity of Xsc (also producing sulfite) into acetyl-CoA, for entry into the TCA and the glyoxylate bypass cycle. We detected the expression of the key enzymes malate dehydrogenase (Mdh) and isocitrate lyase (AceA), from each cycle, respectively, suggesting that the gyoxylate bypass is used for anabolic purposes, while sulfite oxidation is used for energy conservation by ‘*Ca*. Taurinisymbion ianthellae’. In addition to the expressed proteins detected for the processing of taurine-derived sulfur and carbon compounds, we detected the expression of alanine dehydrogenase (Ald), which would replenish the pyruvate needed by the Tpa enzyme for taurine dissimilation (while also providing NADH/H^+^ for respiration) and provide NH3 for assimilation into glutamine, by a highly expressed glutamine synthetase (GlnA; 1.30% NSAF). Ammonia can also be transported out of the cell via an encoded ammonium transporter (*amtB* ; Fig. 3, Table S2), where it can be readily oxidized to nitrite, as previously demonstrated [10], by the dominant thaumarchaeal symbiont. Excluding the expression of glutamine synthetase, the other detected 6 proteins involved in the catabolism of taurine (Tpa, Xsc, AprA, QmoB, and HdrB) amounted to 0.97% NSAF of the 149 proteins assigned to the ‘*Ca.* Taurinisymbion ianthellae’ MAG. Nevertheless, the detection of the high affinity GlnA suggests that ‘*Ca.* Taurinisymbion ianthellae’ was nitrogen limited at the time that the *I. basta* holobiont individual was sampled *in situ*. It has been previously observed that the *I. basta* holobiont, along with its thaumarchaeal ammonia-oxidizing symbiont, ‘*Ca.* Nitrosospongia ianthellae’, is nitrogen limited at ambient conditions[10], and this is in line with apparent nitrogen limitation in marine systems [101] and mammalian large intestines [102]. The observations that the *I. basta* holobiont readily dissimilated added taurine in *I. basta* explant incubations (see below) and incorporated the taurine-derived carbon and nitrogen at C:N ratios indicative of nitrogen limitation (see Supplementary Information), further corroborate these findings.

While the expression of the taurine transporter has been observed in the γ-symbiont of *Olavius algarvensis* [103], the recent observations that marine sponge symbionts have the capacity for taurine utilization, is primarily based on gene presence [25, 26, 35–37], with the exception of metaproteomic evidence for *Ca*. Entotheonella phylotypes residing in the marine sponge *Theonella swinhoei* Y. However, this symbiont expressed multiple encoded copies of TauD-related proteins postulated to possibly function in the breakdown of small aromatic compounds [104]. A recent study by our group performed deep sequencing runs on more *I. basta* individuals — while also conducting metaproteomics — and was able to assemble additional low abundant microbial symbiont MAGs that were previously overlooked [105]. One of these low abundant symbionts is an Alphaprotebacterium (Alphaproteobacteria, g_UBA2767) that also possesses the capability to fully convert taurine into ammonium and sulfate via the same pathways as ‘*Ca*. Taurinisymbion ianthellae’. However, this low abundance Alphaproteobacterium only represents 1–5% of the microbial community and did not express any of the proteins involved in taurine import, taurine dissimilation or sulfite oxidation (it only expressed 0.4% of all detected proteins in the metaproteomic dataset). Moreover, this set of metaproteomic analyses was able to detect the TauABC and AprA proteins and definitively assign them to the ‘*Ca*. Taurinisymbion ianthellae’ MAG.

Oxidation of sulfite to sulfate by taurine-degrading bacteria occurs via different pathways. Frequently periplasmic and cytoplasmic sulfite dehydrogenases are used [106, 107], but these enzymes were not detected in the ‘*Ca*. Taurinisymbion ianthellae’ MAG. The genetic capability to carry out the two-step AMP-dependent indirect oxidation of sulfite to sulfate via APS in the cytoplasm employed by this symbiont, is infrequently detected in taurine degraders, with examples found among the ubiquitous, pelagic SAR11 ([58]; but most SAR11 lack ATP sulfurylase), the γ-3 symbiont in the gutless oligochaete, *Olavius algarvensis* [108], and two marine sponge symbionts [25, 26]. Although a metaproteomic study in Antarctic waters has previously revealed the expression of SAR11-associated peptides for this particular pathway of taurine dissimilation (Tpa, Xsc) followed by sulfite oxidation (AprAB) [58], SAR11 in culture has not yet been shown to excrete sulfate in the presence of taurine [109, 110].

### Taurine metabolism in the Ianthella basta holobiont

Based on the findings of the metaproteogenomic data set, we investigated taurine metabolism in the *I. basta* holobiont experimentally. Firstly, NMR and LC-MS analyses of the 90% MeOH extract and the aqueous Fraction 1 confirmed that taurine was present in this sponge (Fig. 4a, Fig. S3)[111]. The natural concentration of taurine, determined by qNMR, was 5.48 ± 0.69 µmol g^-1^ sponge wet wt, which is in the mid-range of previously reported values for six other marine sponges (1.05–12.5 μmol g^-1^ sponge wet wt sponge)[112]. Congruently, we were able to detect ^13^C and ^15^N enrichment in whole sponge and total nucleic acids elemental analyzer isotope ratio mass spectrometry (EA-IRMS) analyses of *I. basta* sponge explants that underwent additions of an equimolar mixture of ^13^C- and ^15^N-labeled taurine in 48 h incubations (Fig. 4b, Fig. S4). While it cannot be excluded that sorption of labeled taurine contributed to the isotopic enrichment in the whole sponge experiments, the detection of labeled total nucleic acids clearly indicated assimilation and incorporation of labelled taurine into *I. basta* holobiont biomass. Thus, from the whole sponge EA-IRMS data, we were able to estimate uptake rates of taurine-derived C and N assimilated into whole *I. basta* holobiont biomass and obtained values of 1.38 ± 0.3 nmol C min^-1^ g^-1^ sponge wet wt (± SE) and 0.74 ± 0.2 nmol N min^-1^ g^-1^ sponge wet wt (± SE). These *I. basta* holobiont taurine-derived carbon uptake rates correspond to approximately 4–10% of the bulk DOC uptake rates reported for the more diverse microbiome of the marine sponge *Aplysina aerophoba* [17], thereby highlighting the importance of taurine in the *I. basta* holobiont. Moreover, NanoSIMS analyses conducted on dissociated sponge extracellular material revealed accumulation of taurine-derived C and N within regions (Fig. S5 & S6) that were similar in size and shape to typical sponge microbial symbiont cells previously visualized in *I. basta* by transmission electron microscopy [55, 113].

**Figure 4.**
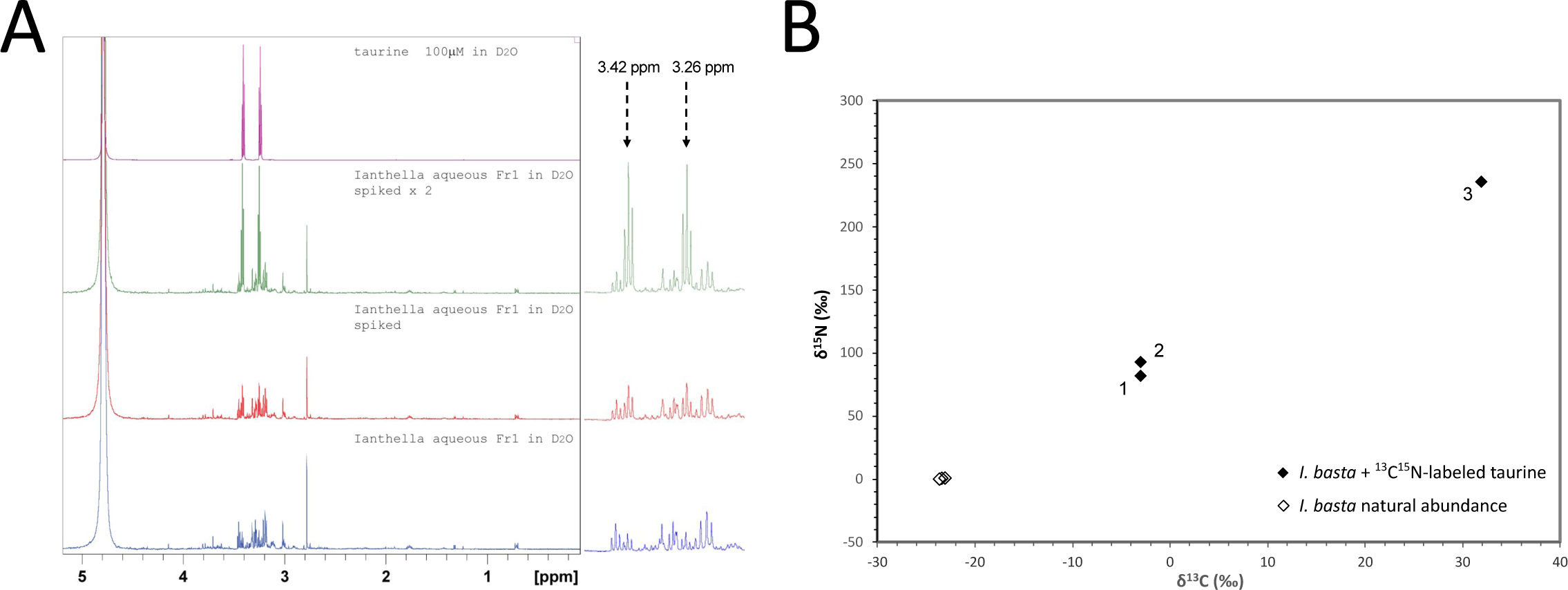
(A) Detection of taurine in the aqueous fraction Fraction 1 of the 90% methanol extract of *Ianthella basta* via ^1^H NMR. ^1^H NMR spectra of Fraction 1 from C18 of crude methanolic extract (blue), Fraction 1 spiked with 32 μl (red) and 128 μl (green) of 100 μM taurine, as well as 100 μM taurine (purple), all acquired in D2O. Zoomed regions are depicted of diagnostic triplet signals at 3.26 and 3.42 ppm, and the increase in taurine signal intensity with spiking. (B) Incorporation of ^13^C- and ^15^N-labeled taurine into sponge holobiont biomass at the total nucleic acid level as determined by EA-IRMS. Data from three sponge explants incubated with labeled taurine (numbered for reference) and three control sponge explants incubated with unlabeled taurine are depicted.

### Taurine catabolism by *Ianthella basta* holobiont results in the accumulation of inorganic S and N species

Some microbial pure cultures catabolizing taurine, oxidize the dissimilated sulfonate moiety to sulfate, which is subsequently excreted [114–116]. Considering that metaproteogenomic analyses predicted sulfate as a secreted end product of ‘*Ca*. Taurinisymbion ianthellae’ taurine metabolism, we analyzed whether the *I. basta* holobiont produced sulfate (SO4) in experiments with and without added taurine. Specifically, we conducted 48 h incubations with *I. basta* explants amended with taurine (in total 1.6 mM) and without taurine in sulfate-free artificial seawater (SFASW). SFASW was used in these incubations to facilitate the detection of sulfate formation by the holobiont. Pairwise comparisons in incubations with sponges in SFASW without the addition of taurine (SFASW + *I. basta*), resulted in a small, but significant production of SO4 after 12 h, indicating *in vivo* production of SO4 (*p* ˂ 0.05, one-tailed t-test) (Fig. 5a). In incubations with *I. basta* explants and added taurine (SFASW + *I. basta* + taurine), we observed a similarly small increase at 12 h (*p* ˂ 0.05, one-tailed t-test), with SO4 production strongly increasing between the 12 h and 48 h time points, suggesting a metabolic lag effect (Fig. 5a). Moreover, the final SO4 concentrations of 590 ± 89 (SE) µM SO4 (59.0 ± 2.0 µmol g sponge wet wt in Fig. 5a) at 48 h were significantly higher than the SO4 concentrations at t = 0 and t = 12 h (one-way ANOVA followed by Holm-Sidak tests, *p* ˂ 0.001) and also significantly higher than all control incubations, including the incubations with sponge explants but without taurine at t = 48 h (one-way ANOVA followed by Holm-Sidak pairwise multiple comparison tests, *p* ˂ 0.001). Two additional incubation experiments, with refined temporal resolution (t = 0, 12, 24, 36 and 48 h) using 1 mM and 100 µM unlabeled taurine amendments, respectively, resembled the results of the previous experiment, and displayed a significant SO4 increase after 48 h for both taurine treatments (one-way ANOVA followed by Tukey pairwise multiple comparison tests, *p* ˂ 0.05), with essentially a full recovery of the +100 µM taurine sulfonate in the form of sulfate^-^ after 48h (Fig. S7).

**Figure 5.**
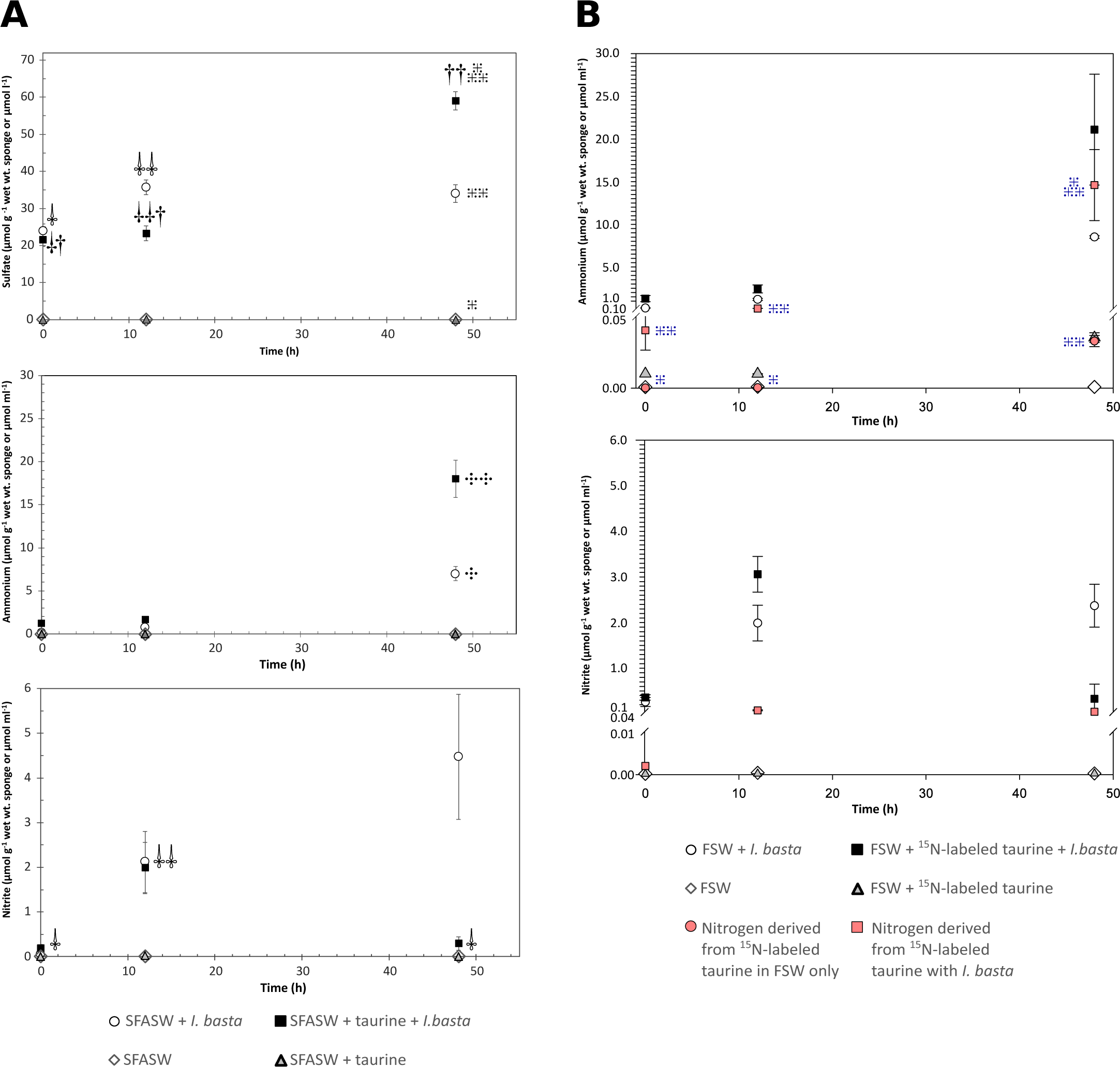
Time series of the concentrations of dissolved sulfate, ammonium, and nitrite in the incubation media of *I. basta* holobiont batch incubations (**A**) 48 h incubation in sulfate-free artificial seawater (SFASW) performed either without (white circle) or with added unlabeled taurine (filled square: t = 0, +1 mM taurine; t = 36 h, +0.6 mM taurine). Concentrations are normalized to the wet weight of the respective *I. basta* explants applied in the incubations (8.83 ± 2.38 SD g sponge wet wt.; range: 6.5–13.4 g sponge wet wt.). The data from the corresponding control experiments with SFASW without *I. basta* holobiont are displayed as white diamonds (without added taurine) and grey triangles (with added taurine) and plotted on the same axes, given in µmol ml^-1^. A starting average concentration of 0.2 mM sulfate was measured in all incubation treatments containing *I. basta* explants most likely representing residual carryover from the sponge material. Incubations were performed in biological triplicates and water samples from each incubation were taken at each time point. In A), ⸸⸸ symbols for sulfate concentrations denote significant difference in pairwise comparisons of incubations from t = 0 (⸸) to t = 12 (⸸⸸) for both the incubations with *I. basta* and taurine and without added taurine (one-tailed t-test, *p* ˂ 0.05); †† symbols in the sulfate concentration panel reflect significant differences for sulfate concentrations for *I. basta* and taurine incubations at t = 48 (††) when compared to the earlier time points (†) (one-way ANOVA followed by Holm-Sidak tests, *p* ˂ 0.001); similarly, the number of 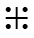 symbols reflects significant differences for sulfate concentrations for incubations with *I. basta* and taurine when compared to all other treatments at t = 48 h (one-way ANOVA followed by Holm-Sidak tests, *p* ˂ 0.001). ⸭⸭ in the ammonium concentration panel reflects a significant difference between incubations with *I. basta* and taurine and only *I. basta* (⸭) (two-tailed t-test, *p* ˂ 0.05). ⸸⸸ in the nitrite concentration panel reflects a significant difference between time points for the incubations with *I. basta* and taurine (one-way ANOVA followed by Tukey pairwise multiple comparison tests, *p* ˂ 0.05). (**B**) Ammonium and nitrite accumulation in *I. basta* holobiont batch incubations (48 h) in natural filtered seawater (FSW) performed either with added taurine (filled symbols: t = 0, +1 mM taurine; t = 36 h, +0.6 mM taurine) or without added taurine (9.45 ± 2.53 SD g sponge wet wt; range: 5.5–12.7 g sponge wet wt). Taurine was added as an equimolar mixture of ^13^C- and ^15^N-labeled taurine to obtain sponge material for subsequent EA-IRMS and NanoSIMS analyses (by sacrificially sampling sponges at the end of the experiment) as well as to measure ^15^N in NH4^+^ and NO2^-^. Incubations were performed in biological triplicates and water samples from each incubation were taken at each time point. The data from the corresponding control experiments with FSW without *I. basta* holobiont are displayed as white circles and diamonds as in A). Pink symbols represent the measured concentrations of ^15^N in ammonium and nitrite that were a product of the added ^15^N-labeled taurine (t = 0, +0.5 mM ^15^N-labeled taurine; t = 36 h, +0.3 mM ^15^N-labeled taurine). The number of ⸭⸭ symbols in the ammonium concentration panel in B) reflects significant differences in ^15^NH4^+^ concentrations measured in incubations with *I. basta* and isotopically labeled taurine and in incubations without *I. basta* between different time points and across treatments (one-way ANOVA followed by Student-Newman-Keuls pairwise multiple comparison tests, *p ˂* 0.05). Error bars refer to +/-1SE.

Furthermore, we observed NH4 and NO2 production in *I. basta* incubations, with and without added unlabeled taurine (SFASW + *I. basta* + taurine; SFASW + *I. basta*), where NH4 production at the end of the experiment with added taurine, drastically exceeded those of *I. basta* incubations without added taurine, consistent with the production of ammonium from taurine (Fig. 5a; *p* ˂ 0.05, two-tailed t-test). Furthermore, a clear increase in NO2 production occurred within the first 12 h and was found to be significant for incubations with added taurine (Fig. 5a, SFASW + *I. basta* + taurine: one-way ANOVA followed by Tukey pairwise multiple comparison tests, *p* ˂ 0.05). NH4^+^ and NO2^-^ production in *I. basta* has previously been observed [10] and the NO2 production was attributed to the only ammonia-oxidizing symbiont, ‘*Ca*. Nitrosospongia ianthellae’. At the end of the experiment, NO2 concentrations decreased in the incubations with added taurine, consistent with the previously demonstrated inhibition of ammonia oxidation at concentrations ≥100 µM NH4 in the *I.* basta holobiont and/or with the concomitant increased hypoxia facilitating denitrification activities [10], a capability that is encoded in ‘*Ca*. Taurinisymbion ianthellae’. Additional experiments with refined temporal resolution, displayed the same general trends (Fig. S7).

In addition, FSW incubations of the *I. basta* holobiont with ^15^N-labeled taurine were performed. In these experiments, we were able to detect ^15^NH4 and NO2 formation by *I. basta* sponge explants (Fig. 5b), demonstrating taurine metabolism and oxidation of taurine-derived ammonia to nitrite by the sponge holobiont. Specifically, this indicates that the dominant thaumarchaeal symbiont hosted by *I. basta*, ‘*Ca*. Nitrosospongia ianthellae’, which was previously shown to be the only resident ammonia-oxidizer in this holobiont [10], oxidized the taurine-derived ammonia to nitrite. Importantly, the amount of ^15^NH4 and NO2 derived from N-labeled taurine in incubations with *I. basta* was always higher than the amount detected in FSW control incubations without *I. basta* at t = 12 and t = 48 h (Figure 5b; one-way ANOVA followed by Student-Newman-Keuls pairwise multiple comparison tests, *p* ˂ 0.05; ^15^NO2 in FSW control incubations were undetectable in all samples). This demonstrates that these successive conversions were catalyzed by holobiont members and that this process starts as early as 12 h, suggesting that the similarly observed increase in SO4 was also derived from taurine. In *I. basta* explant incubations without externally added taurine (FSW + *I. basta*), NH4 and NO2 concentrations still greatly exceeded those incubations without sponge explants and added taurine (FSW + taurine), providing additional support for the production of these compounds by *I. basta* holobiont members. Finally, HPLC analysis of taurine concentrations for these FSW incubations confirmed uptake of taurine by the *I. basta* holobiont, since incubations with *I. basta* explants and added taurine, exhibited concentrations (FSW + *I. basta* + taurine: 1.20 ± 0.05) significantly lower than in control incubations without *I. basta* explants and added taurine (FSW + taurine: 1.73 ± 0.02 mM taurine; Kruskal-Wallis, *p* < 0.05; Fig. S8).

These incubation experiments demonstrate that the *I. basta* holobiont explants actively processed and incorporated taurine-derived carbon and nitrogen, while then dissimilating the resultant ammonia moiety to nitrite and oxidizing the sulfonate moiety to sulfate, from the onset through the course of the incubations. Taken together with the metaproteogenomic dataset, we conclude that ‘*Ca*. Taurinisymbion ianthellae’ is the holobiont member most likely responsible for this taurine incorporation and dissimilation, including the oxidation of the resultant sulfite to sulfate. Although the observed rates are potential net rates under specific incubation conditions, which will undoubtedly differ from *in situ* rates, as in all *ex vivo* experiments, the prompt incorporation and dissimilation of added exogenous taurine, suggests a N-limited sponge holobiont that is metabolically ready to access and transform this compound for anabolic as well as catabolic purposes. Recently, several laboratory and field-based studies have documented the exchange of sulfonates between marine bacteria and eukaryotic phytoplankton, with taurine (or its derivatives), always being one of the main sulfonates involved [32–34]. In co-culture, the coastal diatom *Pseudo-nitzschia multiseries* apparently provides a *Sulfitobacter* strain (SA11) with taurine in exchange for the plant hormone indole-3-acetic acid as well as ammonium [38]. Similarly, cross-feeding between nitrite oxidizers and ammonia oxidizers has been demonstrated wherein urea or cyanate is first processed by nitrite oxidizers [117, 118], thereby providing the cleaved ammonia for oxidation to nitrite. While marine Thaumarchaeota have been shown to directly utilize urea and cyanate [119] as well as taurine exuded by mesozooplankton [120], the mechanisms for the direct utilization of these dissolved organic compounds has yet to be elucidated and indirect utilization remains a confounding factor.

### A proposed model of an auxotrophy network between symbionts and the sponge host

In the *I. basta* metagenome we found no indications for taurine biosynthesis by its microbial symbionts. Consequently, we believe it to be most likely that taurine detected in the *I. basta* holobiont is synthesized by the sponge and after release, via export or sponge cell lysis, is utilized by ‘*Ca*. Taurinisymbion ianthellae’. In fact, the 5.5 µmol g^-1^ sponge wet weight taurine we measured in *I. basta* would correspond to ∼6.6 mM concentrations within the sponge explants used for the incubations, (under the assumption that 1.2 g sponge wet wt = 1 cm^-3^; [96]) which is on the high end of measured cytosolic concentrations in eukaryotic phytoplankton [34], and this still does not take into account further potential concentration in specialized cells, cell compartments, or in the mesohyl. We thus propose that endogenous taurine is either translocated by *I. basta* to ‘*Ca*. Taurinisymbion ianthellae’, as has been shown recently for DOM-derived carbon and nitrogen in a high microbial abundance (HMA) sponge [15], or is derived from host cell lysis, as a result of the rapid cell turnover characteristic of marine sponges [121, 122]. Nevertheless, it is possible that marine sponges themselves process taurine from the environment, as has been shown for bulk DOM mixtures [15, 17], instead of synthesizing it, and there remains the possibility that in our incubation experiments *I. basta* itself took up taurine and processed it. Even though our incubation experiments involved the addition of exogenous taurine, the almost immediate and continued production of the dissimilated and oxidized end products, ammonia and sulfate, two processes for which, so far, only microbially mediated pathways have been demonstrated, suggests that the added taurine was processed primarily by the dominant gammaproteobacterial symbiont, ‘*Ca*. Taurinisymbion ianthellae’, and that it is metabolically ‘primed’ to process this compound. Alongside the incubation experimental results and the measured taurine concentrations in *I. basta*, we therefore assume endogenous taurine supplementation by *I. basta* to be the most parsimonious route given the low taurine seawater concentrations [34, 120], competition with other microbes in seawater where taurine turnover rates are high [120], the ability of the sponge to be able to gain the taurine precursors cysteine and methionine from multiple sources (exogenous DOM and POM, as well as from its microbial symbionts), and the expression of the taurine degradation pathway by ‘*Ca*. Taurinisymbion ianthellae’ *in vivo*.

Taurine biosynthesis is typically considered to be a metazoan feature but has not yet been demonstrated experimentally in sponges nor has this capability been suggested in sequenced sponges. More recent discoveries have demonstrated bacterial and microalgal capabilities in synthesizing taurine [123, 124], and yet taurine biosynthesis in microalgae and bacteria is not universal and quite sporadic in distribution [34, 125]. Taurine biosynthesis pathways in mammalian and invertebrate tissues use cysteine as a precursor [45] and references therein] and cysteine synthesis in metazoans requires methionine [126]. The methionine synthesis pathway, along with the biosynthesis pathways for several other essential amino acids, are generally lacking in metazoans [127–129], and thus these compounds are typically acquired via heterotrophy. While at least some sponges seem to possess the genomic repertoire for cysteine and methionine synthesis [130, 131], so far, no studies have confirmed their biosynthesis in radiolabeled amino acid precursor or feeding studies, whereas scleratinian corals can produce low amounts of methionine and perhaps even cysteine [131, 132]. ‘*Ca*. Taurinisymbion ianthellae’ encodes pathways for the synthesis of methionine and cysteine (Fig. S9), which are connected with the degradation of taurine and DMSP (Fig. 3), and also encodes several copies of EamA domain-containing proteins (IPR000620) and two copies of LeuE-type export proteins (IPR001123) (Fig. 3 & Table S5), with both families containing homologous representatives demonstrated to export cysteine [133] and methionine [134], respectively. Although further studies are needed to confirm whether marine sponges can indeed synthesize cysteine and methionine (as well as taurine), it is tempting to speculate that ‘*Ca*. Taurinisymbion ianthellae’ synthesizes and provides methionine and cysteine, either via active transport or symbiont cell lysis, to *I. basta*, and that the latter is used as a precursor for taurine biosynthesis. Cysteine or methionine auxotrophy in *I. basta* is not a precondition for this metabolic exchange to occur, as highly regulated kinetic switches may exist within the *I. basta* holobiont that is dependent on dynamic shifts in resource limitation and environmental stressors.

Although the methionine synthase (MetH)-containing pathway for methionine biosynthesis in ‘*Ca*. Taurinisymbion ianthellae’ is vitamin B12 dependent (this organism also encodes a vitamin B12-independent pathway using DMSP as precursor as well as the cobalamin-independent methionine synthase, MetE; Fig 3, Fig S9), its MAG does not contain the genes required for synthesis of this vitamin, while it does encode a putative vitamin B12 transporter BtuCDF (Fig. 3 & Table S5). As the thaumarchaeal symbiont of *I. basta* ‘*Ca.* Nitrosospongia ianthellae’ has the genomic repertoire for cobalamin (vitamin B12) biosynthesis [10] it is conceivable that it provides vitamin B12 for methionine biosynthesis in ‘*Ca*. Taurinisymbion ianthellae’ if DMSP is not available, while receiving ammonia produced by the γ-symbiont from taurine degradation. Nitrite formed by ammonia oxidation by ‘*Ca.* Nitrosospongia ianthellae’ can then serve under low oxygen concentrations as electron acceptor in ‘*Ca*. Taurinisymbion ianthellae’. Furthermore, since nitric oxide has been shown to have an immediate contractile effect on sponges [135] and seems to be an ancient and key regulator in the marine sponge life cycle [136], the fact that ‘*Ca*. Taurinisymbion ianthellae’ encodes two denitrification genes — a nitrite reductase, *nirK* and a single subunit quinol-dependent nitric oxide reductase, *qNOR* [137] — that produces and consumes NO, suggests that this symbiont may be able to modulate host behavior. The frequent presence of an incomplete denitrification pathway along with the quinol-dependent *qNOR* — a nitrite reductase commonly found in pathogenic species [138] — in other sponge metagenomes [28], further points at the possibility that this is a widespread feature of microbial symbionts residing in marine sponges. Taken together, our data indicate a tightly connected metabolic network in the *I. basta* holobiont (Fig. 6), albeit in contrast to selected other sponges, no data are yet available for *I. basta* on the uptake of amino acids directly from the environment [16, 17] and on the acquisition of amino acids from ingested free-living and symbiotic prokaryotes [139].

**Figure 6.**
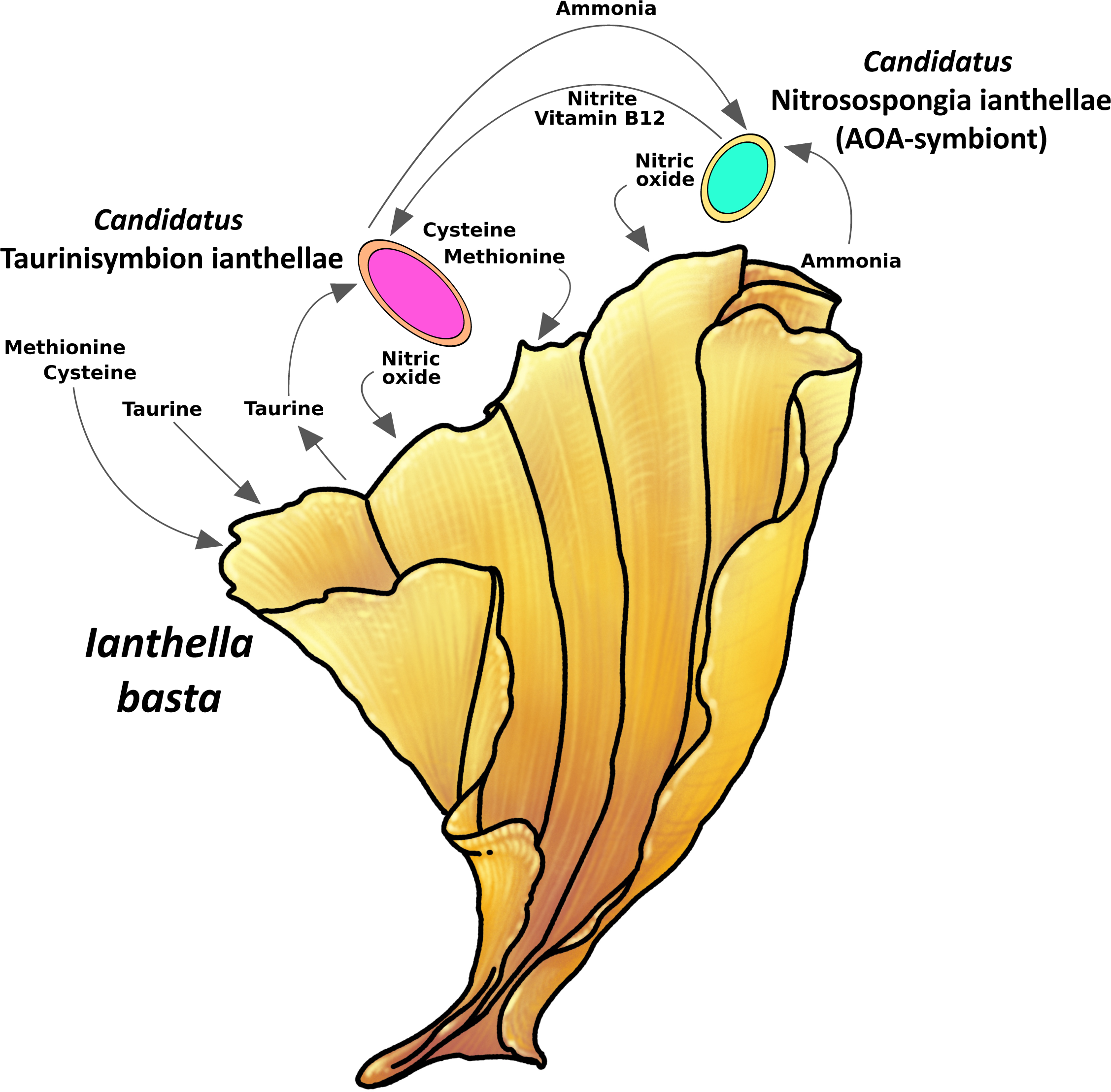
scheme of tripartite metabolic interactions within the *Ianthella basta* holobiont.

## Conclusions

In this study, we showed through a combination of metaproteogenomic analyses and incubation experiments combined with isotope labeling, that the dominant *I. basta* gammaproteobacterial symbiont, ‘*Ca*. Taurinisymbion ianthellae’, most likely uses endogenously occurring taurine, that is potentially produced by the sponge holobiont, for assimilation and energy conservation. We experimentally demonstrated that the taurine dissimilation products, sulfate and ammonia, are secreted and that the latter compound is used for ammonia oxidation by the dominant thaumarchaeal symbiont — ‘*Ca.* Nitrosospongia ianthellae’. Taurine seems to be an important metabolite, not only in *I. basta*, but in many sponge holobiont systems, as indicated by the seeming ubiquity of taurine and taurine dissimilating genes within sponge microbiomes [25, 26, 35–37], and the appearance of a specific taurine-dissimilation/sulfite-oxidation pathway restricted mostly to symbiotic microorganisms [25, 26, 108]. Taurine also appears to be a central metabolite in several other metazoan holobiont systems such as in cnidarians, various bivalve species [44, 45, 140, 141], and in mammalian gut systems [39, 40] . Future research is needed to further disentangle the mechanisms of taurine synthesis via symbiont-derived or environmentally supplied precursors in marine sponge holobionts, to better understand the role of exogenous and endogenous taurine uptake, and the numerous roles taurine may play for the functioning of these symbioses.

## Acknowledgements

We thank Emmanuelle Botte for assisting with experimental incubations, Heidi Luter for assisting with field collections and Bettina Glasl and Pam Engelberts for helpful discussions. Florian Moeller and Maria Mooshammer were supported by the ERC Advanced Grant Nitricare (294343 to MW). We thank Bernhard Schink for valuable advice regarding the provisional naming of the gammaproteobacterial symbiont.

