## Supplementary Information for "Taurine as a key intermediate for host-symbiont interaction in the tropical sponge *Ianthella basta*"

**Table of contents:**

1. Supplementary Materials and Methods
2. Supplementary Results and Discussion
3. Figures S1 – S9
4. Table S1
5. Supplementary Note
6. References

### Supplementary Materials and Methods

***NanoSIMS analysis of* Ianthella basta**. NanoSIMS measurements were performed on a NanoSIMS 50L (Cameca, Gennevilliers, France) on two of the three *I. basta* explant samples obtained from the $^{13}C^{15}N$-taurine incubations at the Large-Instrument Facility for Environmental and Isotope Mass Spectrometry at the University of Vienna. Specimens were selected for maximum isotope enrichment, based on results from EA-IRMS bulk analysis (0.62 and 0.55 at% $^{15}N$; 1.19 and 1.15 at% $^{13}C$). Prior to data acquisition, analysis areas were pre-sputtered utilizing a high-intensity, slightly defocused $Cs^+$ ion beam (400 pA beam current, ~1 µm spot size) until a $Cs^+$ fluence of 3.2E15 ions $cm^{-2}$ was obtained. Data were subsequently acquired as multilayer image stacks by sequential scanning of a finely focused $Cs^+$ primary ion beam (*ca*. 80 nm probe size at 2 pA beam current) over areas between 10.3 × 15 and 29 × 29 $\mu m^2$ at 176 × 256 and 512 × 512 pixel image resolution and a primary ion beam dwell time of 10 to 40 msec (pixel*cycle)$^{-1}$ , corresponding to a per-cycle $Cs^+$ fluence in the range from 8.0E13 to 1.5E16 ions $cm^{-2}$ per cycle. $^{12}C^-$, $^{13}C^-$, $^{12}C^{12}C^-$, $^{12}C^{13}C^-$, $^{12}C^{14}N^-$, $^{12}C^{15}N^-$, $^{31}P^-$ or $^{32}S^-$, as well as secondary electrons, were detected simultaneously. Signal intensities were corrected for detector dead time and quasi-simultaneous arrival (QSA) effects using QSA sensitivity factors (i.e., “beta” values) of 1.1, 1.1, 1.06, 1.06, 1.05, 1.05, 1.1, 1.1 for $^{12}C^-$, $^{13}C^-$, $^{12}C^{12}C^-$, $^{12}C^{13}C^-$, $^{12}C^{14}N^-$, $^{12}C^{15}N^-$, $^{31}P^-$, and $^{32}S$, respectively.

Images based on NanoSIMS measurement data were generated using the Open MIMS plugin [1] in the FIJI package based on ImageJ [2]. C isotope composition images displaying the $^{13}C/(^{12}C+^{13}C)$ isotope fraction, designated as atom% $^{13}C$, were inferred from the $C_2^-$ secondary ion signal intensity distribution images via per-pixel calculation of $^{13}C^{12}C^-/(2\cdot{}^{12}C^{12}C^-+^{13}C^{12}C^-)$ intensity ratios, while N isotope composition images displaying the $^{15}N/(^{14}N+^{15}N)$ isotope fraction, designated as atom% $^{15}N$, were inferred from the $^{12}CN^-$ secondary ion signal intensity maps via per-pixel calculation of $^{12}C^{15}N^-/(^{12}C^{15}N^-+^{12}C^{14}N^-)$ intensity ratios. Secondary electron images were utilized for representation of the sample morphology.

Region of interest (ROI) specific numerical data evaluation was also conducted using the Open MIMS plugin. ROIs were defined according to the isotope enrichment patterns since it was not possible to unambiguously identify characteristic cellular features. This was likely because analysis areas were accessed via depth-profiling, which is less suited for visualization of cellular (ultra)structure than e.g., cross-sectional analysis of resin embedded samples. C and N isotope compositions were calculated from the accumulated intensities of $^{12}C^{12}C^-$ and $^{12}C^{13}C^-$ and $^{12}C^{14}N^-$ and $^{12}C^{15}N^-$ secondary ion signals detected within each ROI. The isotopic composition values shown in Fig. S6 were determined via stepwise inspection of the isotopic enrichment patterns displayed in each individual image of the acquired multilayer stacks (Fig. S7). As such, the displayed values may rather be considered as conservative estimates of the isotopic enrichment due to carry-over (e.g., through re-deposition and/or atomic mixing) of atoms and molecules from less enriched regions within the sponge extracellular matrix during the measurement process (including pre-sputtering). The analytical uncertainty of the isotope composition values (indicated by the error bars in Fig. S6), emerging from the random error in single ion counting, was estimated on the basis of Poisson statistics and calculated from the signal intensities (given in total counts within individual ROIs) via:

atom% $^{13}C$ inferred from $C_2^-$ signal intensities:

$$\sigma_{Pois,\ at\%13C}=100/(2\cdot{}^{12}C^{12}C^- + {}^{12}C^{13}C^-)^2 \cdot sqrt(({}^{12}C^{12}C^-)^2 \cdot {}^{12}C^{13}C^- + ({}^{12}C^{13}C^-)^2 \cdot {}^{12}C^{12}C^-)$$

atom% $^{15}N$ inferred from $CN^-$ signal intensities:

$$\sigma_{Pois,\ at\%15N}=100/({}^{12}C^{14}N^- + {}^{12}C^{15}N^-)^2 \cdot sqrt(({}^{12}C^{14}N^-)^2 \cdot {}^{12}C^{15}N^- + ({}^{12}C^{15}N^-)^2 \cdot {}^{12}C^{14}N^-)$$

### Supplementary Results and Discussion

**Dimethylsulfoniopropionate (DMSP) catabolism and other notable metabolic features.** The main sources of DMSP are micro and macro-algae [3], for which it serves as an organic osmolyte. As DMSP is ubiquitous in marine surface waters and is detected in concentration from less than 1 nM in the open oceans to several micromolar in phytoplankton blooms, this compound will also be available for filter-feeding sponge symbionts. DMSP is a source of reduced sulfur and carbon for sponge symbionts microbes and the precursor for the microbially-mediated formation of the climatically active gas dimethylsulfide (DMS). Within the '*Candidatus* Taurinisymbion ianthellae' MAG, we detected an ABC transporter (*OpuABC*) and a betaine-choline-carnitine-transporter (BCCT; *RbtB*/*SbtA*) that have both been shown to import DMSP [4, 5] along with the key genes for the DMSP cleavage (*dddP*) and demethylation (*dmdA*) pathways (Figure 3). The encoded enzymes DddP and DmdA catalyze conversion of DMSP to DMS and acrylate and to 5-methyltetrahydrofolic acid (5-Methyl-THF) and methylmercapto-propionate (MMPA), respectively. Both enzymes are abundant in marine microorganisms [6]. Although the DMSP cleavage pathway has been found in Gammaproteobacteria [7, 8], in general this pathway is more frequently found in Alphaproteobacteria [7, 9]. In addition to the DMSP cleavage and demethylation genes, we were able to identify and detect the expression of many of the respective downstream enzymes that are used to assimilate or catabolize the initial intermediates of both pathways (Table S4). For the DMSP demethylation pathway, we found homologous genes necessary for MMPA degradation to methanethiol (*dmdB*, *dmdC*, and *acuH*) [9, 10] as well as for the removal of the methyl group from DMSP. Although we did not detect DmdA or DddP expressed in the metaproteomic dataset, we were able to observe the expression of the OpuC transporter subunit, and for the DMSP cleavage pathway, the expression of most of the enzymes necessary for the degradation of acrylate to the coenzyme A (CoA) derivatives propionyl-CoA (PrpE, AcuI) and acetyl-CoA (AcuH, DddC; Figure 3) [9] with the exception of the DmdD enzyme. The AcuH enzyme has been shown to exhibit activity on acrylate/acryloyl-CoA as well as on the DmdD substrate methylthioacrylyl-CoA (MTA-CoA) [9] (and references therein), thereby allowing it to function in both the DMSP cleavage and demethylation pathways. The hydration of MTA-CoA leads to the production of methanethiol, acetaldehyde, $CO_2$, and free CoA [11], and our detection of three genes in the cystathionine beta-lyases/cystathionine gamma-synthases family (*metB*, *metZ*, *metY*), two of which have been shown to convert methanethiol to methionine [12, 13], suggests the assimilation of sulfur derived from DMSP into the γ-symbiont as methionine (Figure 3 and S9). These family of enzymes are pyridoxal phosphate (PLP) dependent, and '*Candidatus* Taurinisymbion ianthellae' has the genetic capacity to produce PLP, thereby providing it with another vitamin $B_{12}$-independent pathway for methionine biosynthesis if DMSP is available (Figure 3 and S9). Finally, '*Candidatus* Taurinisymbion ianthellae' has the metabolic potential to use the 5-methyl-THF produced by DMSP demethylation (i) as a methyl donor for methionine, (ii) as a carbon donor in the serine assimilation pathway after oxidation into 5,10-methylene-THF by an encoded *metF*, or (iii) a as source of ATP and reductant when 5,10-methylene-THF is oxidized to formate and then to $CO_2$. In the metaproteome, we detected enzymes for C-1 assimilation (GlyA, Hpr, Mdh, and Mcl) as well as for 5,10-methylene-THF oxidation to formate (Fhs) (Figure 3; Tables S3 and S4).

DMSP catabolizing genes have been intermittently found in microbiome members of marine sponges [14–16]. While no data for *Ianthella basta* are available, DMSP concentrations in some other marine sponges were found to be relatively low or undetectable (3 of 5 sponges: 0.2–3 µmol/g wet wt.) [17] while zooxanthellae-harboring invertebrates such as corals and anemones are known to contain much higher concentrations of this compound [17, 18]. Similarly, phytoplankton, marine plants and more recently marine bacteria have been shown to synthesize DMSP (reviewed in [19]), but we did not detect either of the two key genes (*dysB* and MSMT) necessary for bacterial DMSP production in the *I. basta*-associated microbial MAGs. Recent single-cell measurements of the model Alphaproteobacterium *Ruegeria pomeroyi* DSS-3 under variable DMSP concentrations [20] demonstrated that at least some marine bacteria typically residing in low mean seawater concentrations of DMSP (16.91 ± 22.17 nM) [21] most likely express both DMSP degradation pathways at baseline levels, upregulate both pathways at 1 µM DMSP, and are thus primed for DMSP maxima either at the spatial microscale or during periodic hotspot events. As marine sponges have been found to be important sinks for DOM in coral reef environments [22] and coral mucus can be assimilated into marine sponge biomass [23], it is conceivable that DMSP from the surrounding ocean environment may be processed by advantageously located marine sponge microbiomes and that benefit from cyclical phytoplankton bloom events and coral mucus release (suggested to follow lunar periodicity) [24]. In this context, it is interesting to note that a cultured representative of the ubiquitous open ocean SAR11 Alphaproteobacteria, also simultaneously express the cleavage and demethylation pathways, and that a kinetic switch dependent on DMSP concentrations may regulate these metabolic processes [11].

Since DMS is a by-product of the first step of the DMSP cleavage pathway, we checked for the presence of a potential DMS-oxidizing protein within the *I. basta*-associated microbial MAGs and found that '*Candidatus* Taurinisymbion ianthellae' encodes a putative trimethylamine (TMA) monooxygenase/flavin-containing monooxygenase with very high similarities (63% AA ID) to two trimethylamine monooxygenases (Tmm) previously demonstrated to co-oxidize DMS at similarly high affinities as TMAs (Table S4) [25, 26]. Although, the oxidation of DMS to dimethylsulfoxide (DMSO) by Tmm, does not result in any net gain of electrons or reducing equivalents, co-oxidation of DMS during periods of TMA depletion may prevent the formation of free radicals and hydrogen peroxide with the subsequent DMSO accumulation potentially supporting further antioxidant activities ([26] and references therein).

In addition to pathways for metabolizing taurine and DMSP, we also identified numerous other genes in the γ-symbiont MAG putatively involved in organosulfur compound degradation, including the dissimilation of sulfopropanediol (*hpsN*), sulfoacetate (*sauS*), sulfolactate (*suyAB*), and cysteate (*cuyA*). Nevertheless, '*Candidatus* Taurinisymbion ianthellae' has the capability to assimilate sulfate via sulfate adenylyltransferase (*sat*), adenylyl sulfate kinase (*cysC*), PAPS reductase (*cysH*), and an assimilatory sulfite reductase (*sir*).

Notably, the '*Candidatus* Taurinisymbion ianthellae' MAG encodes for a wide range of transporters representing different families, but primarily characterized by the preponderance of ATP-binding cassette (ABC) transporters, many of which were found to be expressed in the metaproteomic dataset (Table S3 & S5). Within this class of transporters, most are predicted to be involved in the transport of nitrogen containing organic compounds such as amino acids, oligopeptides, polyamines (spermidine and putrescine), compatible solutes (glycine, betaine, and proline), as well as the vitamins thiamin and cobalamin (B1 and B12).

**Fractions of taurine-derived carbon and nitrogen determined within the holobiont.** The C:N ratio of the assimilated taurine by the *I. basta* holobiont was 1.86 ± 0.09 (SD) and 2.37 ± 0.03 (SD), for the whole sponge and nucleic acids-based analyses, respectively which were lower than the corresponding total C:N ratios of 4.6 ± 0.07 (SD) and 3.21 ± 0.2 (SD) determined in the same sponges. These values were also lower than those found for sponge clones that were not incubated with isotope-labeled taurine (4.6 ± 0.15 and 3.40 ± 0.03). This indicates that the holobiont assimilated the taurine-derived nitrogen more efficiently than the corresponding carbon, which is consistent with the inherent C:N stoichiometry of taurine (i.e., 2.0), and suggests that the *I. basta* was nitrogen limited during the course of the incubations. *Alcaligenes defragans* NKNTAU, when grown in pure culture with taurine as a sole carbon and nitrogen source, respires 50% of taurine-supplied carbon and converts the other 50% into biomass, while it incorporates 20% of taurine-derived nitrogen into biomass. The remaining taurine-derived nitrogen is accumulated in the medium as $NH_4^+$, with $SO_4^{2-}$ also produced in a near 1:1 ratio to the taurine supplied [27]. *A. defragans* NKNTAU was also found to express Tpa, Xsc and a Pta when supplied with taurine, thereby converting taurine into acetyl-CoA [28]. In our taurine incubation experiments, we determined a comparable value of 21.0 ± 9.1 % (SE) of fixed taurine nitrogen in the *I. basta* holobiont biomass, obtained from normalization of the amount assimilated into the holobiont biomass to the total amount of metabolized taurine (comprising the fraction of assimilated nitrogen and the released $NH_4^+$ — see Material and Methods for details). The mean amounts of assimilated nitrogen and released ammonia were 20.74 ± 7.7 (SE) µmol N assimilated and 104.1 ± 6.0 (SE) µmol N holobiont tissue per g holobiont tissue wet wt, respectively. However, these values do not take into account the potential loss of dissolved inorganic nitrogen via denitrification and dissimilatory nitrate reduction to ammonium (DNRA). Similarly, rapid sponge cell turnover may also result in the loss of fixed N biomass.

By assuming that the amount of $^{15}N$ released as ammonium and nitrite or fixed into sponge biomass is indicative of the total taurine processed, the relative amount of $^{13}C$-carbon fixed would represent 40.1 ± 19 % (SE; range of 77.3–20.0 %) of the supplied taurine $^{13}C$-carbon in the incubation experiments. In concordance with the above interpretation that the *I. basta* holobiont is N-limited, this would additionally indicate that the amount of taurine supplied was in excess of the carbon biomass requirements of the *I. basta* holobiont, and that the rapid choanocyte turnover characteristic of marine sponges [29, 30] may have contributed to the additional loss of supplied carbon. The incubations with isotopically labeled taurine were conducted in natural seawater and due to its high $SO_4^{2-}$ concentration no reasonable $SO_4^{2-}$ flux estimations for inferring taurine consumption could be made. In incubations conducted in SFASW with unlabeled taurine, we observed near stoichiometric production of $SO_4^{2-}$ from the added 100 µM taurine after 48 h, while net $SO_4^{2-}$ recovery in incubations with 1 mM and 1.6 mM taurine conducted in SFASW were 350 ± 18 and 286 ± 36 (SE) µM $SO_4^{2-}$ after 48 h, respectively. Using the net flux of $SO_4^{2-}$ derived from the 1.6 mM taurine incubations in SFASW as a measure of the total amount of taurine metabolized by the sponge holobiont would then yield inferred carbon and nitrogen incorporation values from taurine of 15.5 ± 8.0 % and 8.1 ± 0.4 % (SE), respectively. Since we demonstrated that 100 µM $NH_4^+$ does not affect $SO_4^{2-}$ flux in the same manner as it does with ammonia oxidation within the *I. basta* holobiont [31], this would indicate that either sulfite oxidation is decoupled from carbon and nitrogen incorporation into biomass, or that the potential physiological stress caused by the high $NH_4^+$ concentrations caused a further increase in the turnover of marine sponge choanocytes.

It is tempting to speculate that the microbial-sized cells exhibiting $^{13}C$- and $^{15}N$- enrichments in the NanoSIMS analyses primarily comprise the '*Candidatus* Taurinisymbion ianthellae' population residing in *I. basta* based on (i) the detection of the key genes for the taurine dissimilation–sulfite oxidation pathway in the '*Candidatus* Taurinisymbion ianthellae' MAG, (ii) the detection of some of the respective proteins in the metaproteome, and (iii) the marked differences in the accumulation of sulfate in incubations with added taurine, which indicate the intracellular oxidation of sulfite, a process for which no other of the dominant microbial members in the *I. basta* holobiont have the apparent capability to mediate. These results are therefore consistent with the recent NanoSIMS-based analyses of marine sponge DOM uptake [32, 33], which demonstrated that high-microbial abundance (HMA) and low-microbial abundance (LMA) sponges have the capability to take up significant proportions of supplied DOM and furthermore, that within one HMA sponge, the symbiotic bacteria are primarily responsible for the processing of externally sourced DOM [33]. Although we added taurine in our incubations as an exogenous source, we also detected sulfate accumulation in the incubation media of all incubations containing *I. basta* without added taurine. In this context, it is important to note that the '*Candidatus* Taurinisymbion ianthellae' MAG, also contains genes for the catabolism of the $C_3$-sulfonates sulfopropanediol (*hpsN*), sulfoacetate (*sauS*), sulfolactate (*suyAB*), and cysteate (*cuyA*), which all can lead to the production of sulfate [34–36] yet these compounds are all primarily derived from eukaryotic phytoplankton and cyanobacteria in the marine environment [36]. In addition to the fact that all incubations were conducted in the dark, cyanobacterial or eukaryotic photosymbionts do not reside in *I. basta* and the $C_3$-sulfonate catabolism genes were not found as expressed as proteins in the metaproteomic dataset. The sulfate concentration enhancement ranging from 109 to 10 µM $SO_4^{2-}$ in *I. basta* incubations without added taurine and conducted in SFASW therefore most likely reflect the processing of endogenously produced taurine.

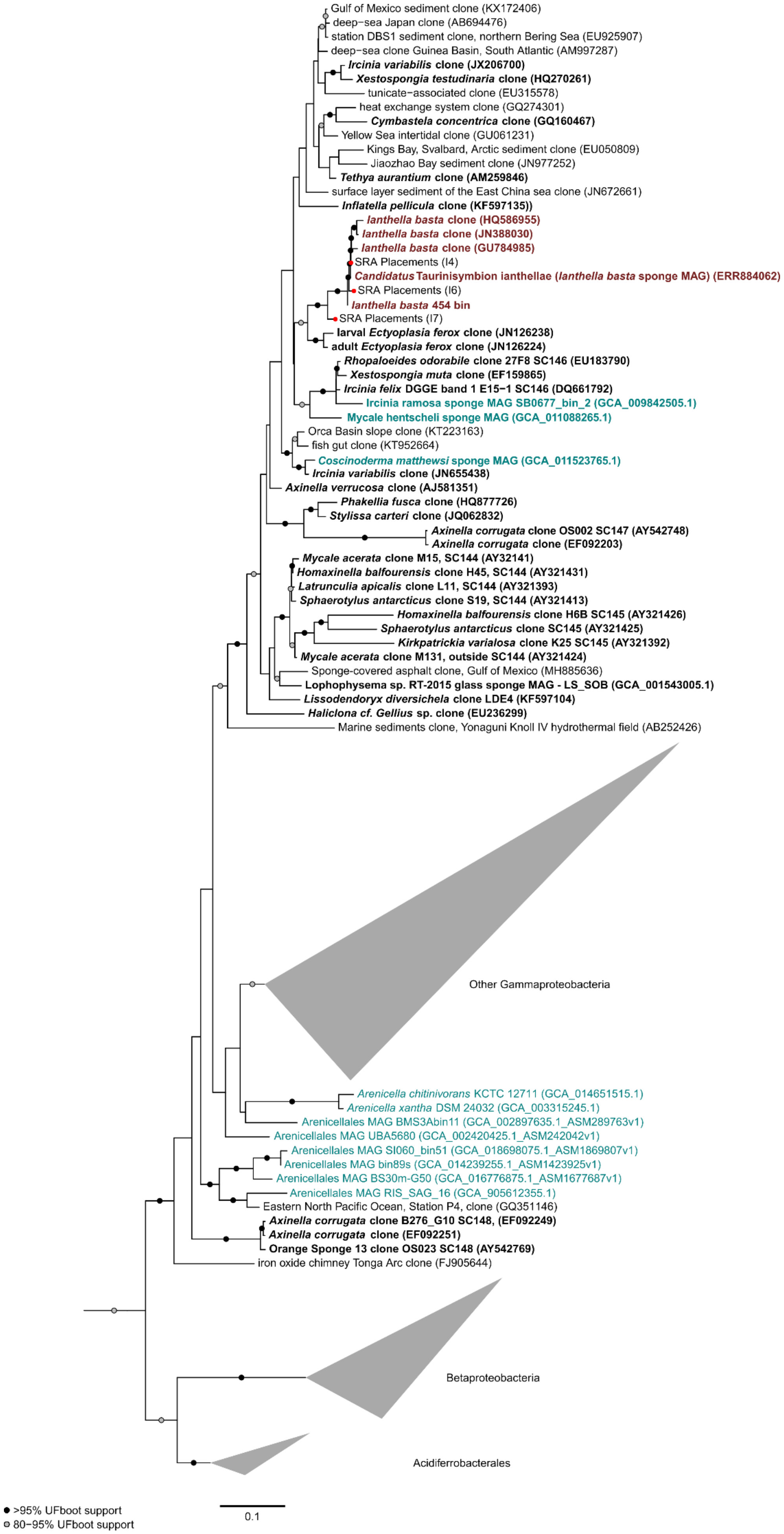

Gulf of Mexico sediment clone (KX172406)
deep−sea Japan clone (AB694476)
station DBS1 sediment clone, northern Bering Sea (EU925907)
deep−sea clone Guinea Basin, South Atlantic (AM997287)
Ircinia variabilis clone (JX206700)
Xestospongia testudinaria clone (HQ270261)
tunicate−associated clone (EU315578)
heat exchange system clone (GQ274301)
Cymbastela concentrica clone (GQ160467)
Yellow Sea intertidal clone (GU061231)
Kings Bay, Svalbard, Arctic sediment clone (EU050809)
Jiaozhao Bay sediment clone (JN977252)
Tethya aurantium clone (AM259846)
surface layer sediment of the East China sea clone (JN672661)
Inflatella pellicula clone (KF597135))
Ianthella basta clone (HQ586955)
Ianthella basta clone (JN388030)
Ianthella basta clone (GU784985)
SRA Placements (I4)
Candidatus Taurinisymbion ianthellae (Ianthella basta sponge MAG) (ERR884062)
SRA Placements (I6)
Ianthella basta 454 bin
SRA Placements (I7)
larval Ectyoplasia ferox clone (JN126238)
adult Ectyoplasia ferox clone (JN126224)
Rhopaloeides odorabile clone 27F8 SC146 (EU183790)
Xestospongia muta clone (EF159865)
Ircinia felix DGGE band 1 E15−1 SC146 (DQ661792)
Ircinia ramosa sponge MAG SB0677_bin_2 (GCA_009842505.1)
Mycale hentscheli sponge MAG (GCA_011088265.1)
Orca Basin slope clone (KT223163)
fish gut clone (KT952664)
Coscinoderma matthewsi sponge MAG (GCA_011523765.1)
Ircinia variabilis clone (JN655438)
Axinella verrucosa clone (AJ581351)
Phakellia fusca clone (HQ877726)
Stylissa carteri clone (JQ062832)
Axinella corrugata clone OS002 SC147 (AY542748)
Axinella corrugata clone (EF092203)
Mycale acerata clone M15, SC144 (AY32141)
Homaxinella balfourensis clone H45, SC144 (AY321431)
Latrunculia apicalis clone L11, SC144 (AY321393)
Sphaerotylus antarcticus clone S19, SC144 (AY321413)
Homaxinella balfourensis clone H6B SC145 (AY321426)
Sphaerotylus antarcticus clone SC145 (AY321425)
Kirkpatrickia varialosa clone K25 SC145 (AY321392)
Mycale acerata clone M131, outside SC144 (AY321424)
Sponge-covered asphalt clone, Gulf of Mexico (MH885636)
Lophophysema sp. RT-2015 glass sponge MAG - LS_SOB (GCA_001543005.1)
Lissodendoryx diversichela clone LDE4 (KF597104)
Haliclona cf. Gellius sp. clone (EU236299)
Marine sediments clone, Yonaguni Knoll IV hydrothermal field (AB252426)
Other Gammaproteobacteria
Arenicella chitinivorans KCTC 12711 (GCA_014651515.1)
Arenicella xantha DSM 24032 (GCA_003315245.1)
Arenicellales MAG BMS3Abin11 (GCA_002897635.1_ASM289763v1)
Arenicellales MAG UBA5680 (GCA_002420425.1_ASM242042v1)
Arenicellales MAG SI060_bin51 (GCA_018698075.1_ASM1869807v1)
Arenicellales MAG bin89s (GCA_014239255.1_ASM1423925v1)
Arenicellales MAG BS30m-G50 (GCA_016776875.1_ASM1677687v1)
Arenicellales MAG RIS_SAG_16 (GCA_905612355.1)
Eastern North Pacific Ocean, Station P4, clone (GQ351146)
Axinella corrugata clone B276_G10 SC148, (EF092249)
Axinella corrugata clone (EF092251)
Orange Sponge 13 clone OS023 SC148 (AY542769)
iron oxide chimney Tonga Arc clone (FJ905644)
Betaproteobacteria
Acidiferrobacterales
>95% UFboot support
80−95% UFboot support
0.1

**Figure S1.** Maximum-likelihood phylogenetic tree of the 16S rRNA gene (after automatic model selection with IQ-Tree) of 'Candidatus Taurisymbion ianthellae'(red and bold) along with other selected sponge (bold font). Entries highlighted in teal represent 16S rRNA sequences extracted from MAGs represented in Figure 1. 'Candidatus Taurisymbion ianthellae' 16S rRNA gene sequences were queried against the SRA database, and the hits were placed into the reference 16S rRNA gene tree (represented on the red nodes) using the Evolutionary Placement Algorithm [37]. Members of the sponge specific clusters (SC) [38] 144-148 are indicated in the tree.

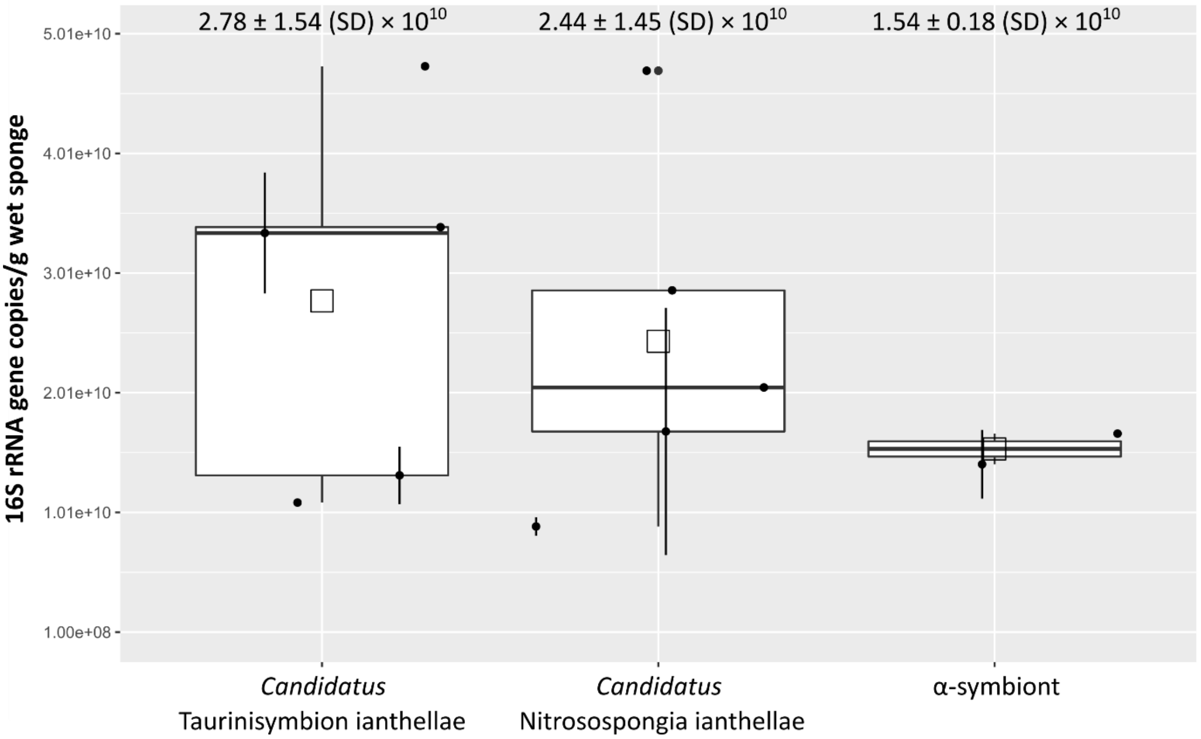


**Figure S2.** Absolute abundance of the three dominant microbial symbionts residing in *Ianthella basta* as quantified using qPCR from five sponge individuals with the exception of the α-symbiont, including those used for metagenome sequencing. Boxplots illustrate summary statistics, indicating the median (horizontal bars), first and third quartile (vertical bars: Q1/Q3 + 1.5 × IQR) as well as the mean (open square). The error bars at the individual data points refer to standard deviation values where samples were measured multiple times. The mean values are also displayed at the top for each microbial symbiont. Based on previously acquired FISH and TEM images from *I. basta* ([39, 40]) we conclude that '*Ca*. Taurinisymbion ianthellae' are most likely rod-shaped cells that are 0.8–1.1 µm long and 0.1–0.28 µm in width. From these assumptions, we can make a calculation of the volume of cells that the estimated abundance of '*Ca*. Taurinisymbion ianthellae' at 2.78 ± 1.5 (SD) x $10^{10}$ 16S rRNA gene copies per g wet sponge, would roughly occupy per $cm^3$. The putative '*Ca*. Taurinisymbion ianthellae' cells would thus occupy ~0.02–0.19% of 1 $cm^3$. Keeping in mind that HMA sponges are considered to have microbial densities of $10^8$–$10^{10}$ per gram of sponge wet weight [41] the abundance numbers of all of the three dominant microbial symbionts within *I. basta* would still add up to $10^{10}$ per gram of sponge wet weight.

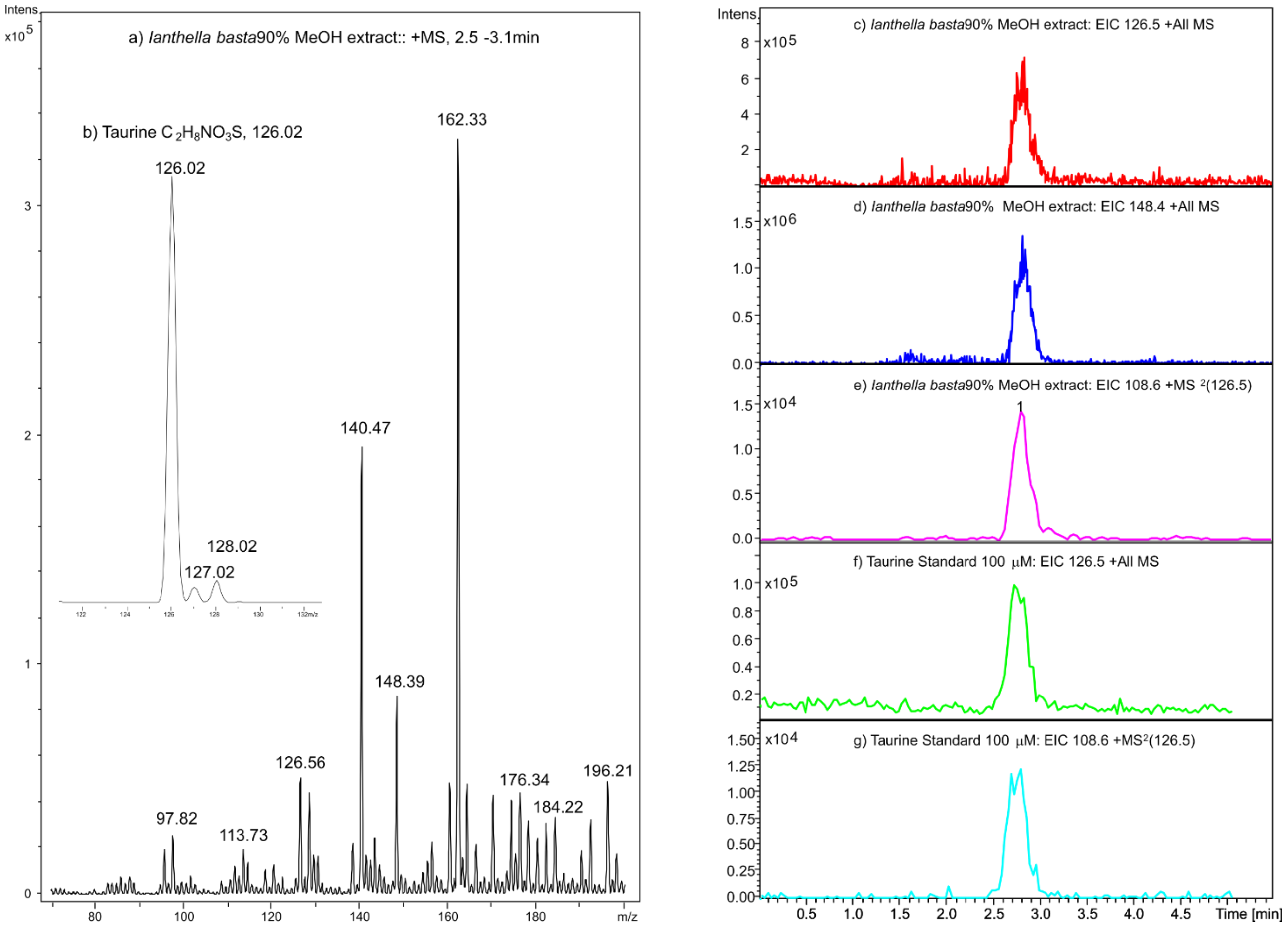


**Figure S3.** LCMS analysis of taurine in *Ianthella basta*. a) Mass spectrum of peak at retention time 2.5–3.1 min for the 90% methanol extract of *Ianthella basta*. Signal at 126.5 *m/z* matches that expected for taurine $[M+H]^+$ ion ($C_2H_7NO_3S$); b) inset shows the expected isotope distribution. Retention time and extracted ion chromatogram (EIC) for c) $[M+H]^+$ (126.5 *m/z*) and d) $[M+Na]^+$ (148.4 *m/z*) of the 90% methanol extract of *I. basta* and e) the expected $[M+H\text{-}H_2O]^+$ fragmentation ion at 108.6 *m/z*, as confirmed by comparison with f) the $[M+H]^+$ and g) $MS^2$ EIC of the taurine standard.

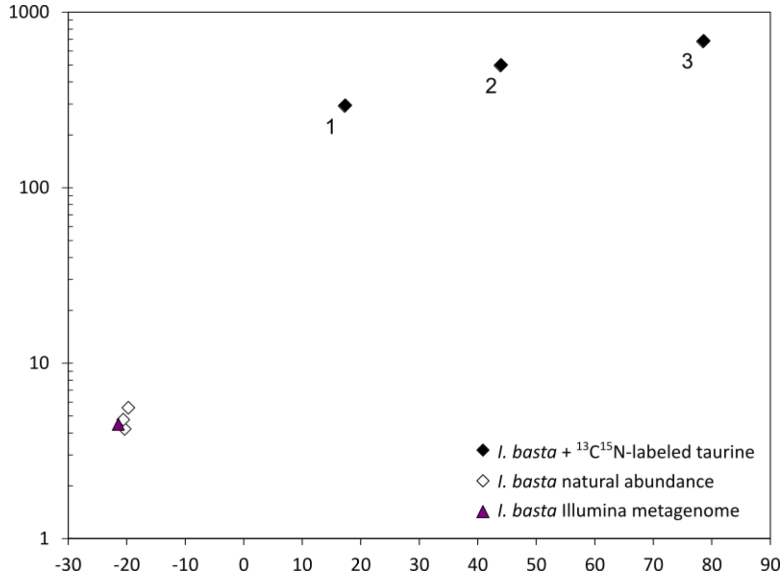


**Figure S4.** $\delta^{13}C$ and $\delta^{15}N$ (‰) values determined in *Ianthella basta* whole sponge tissue homogenate. Three biological replicates (numbered) were analyzed after incubation with isotopically labeled taurine and unlabeled taurine (natural abundance control), respectively. In addition, total nucleic acids (Fig. 4) were extracted from fractions of the samples, for which the results for whole sponge tissue homogenate are depicted. In the analysis, the sponge sample that was used for Illumina-sequencing based metagenomic analysis was also included as an additional natural abundance control.

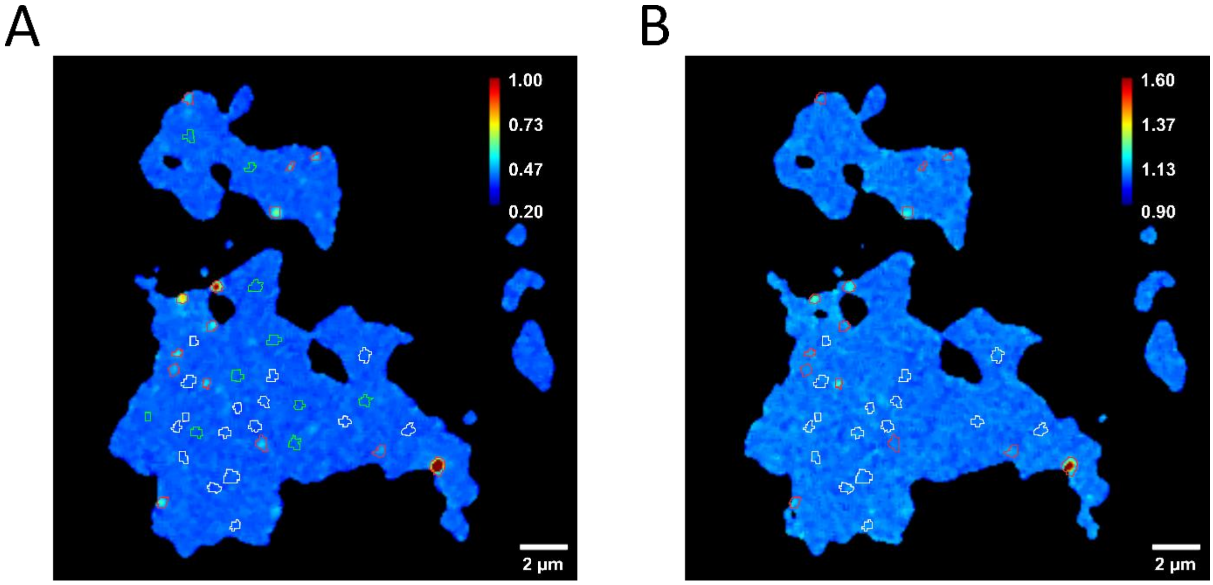


**Fig. S5.** Illustration of the definition and classification of regions of interest (ROI) for numerical evaluation of NanoSIMS measurement data in a representative image from a particular acquisition cycle from *I. basta* extruded extracellular material with respect to the local $^{15}N$ (A) and $^{13}C$ (B) contents (c.f. Fig. S6). The displayed images were acquired at an erosion depth below the surface of the sample, visualizing the presence of isotopically enriched compounds within the sponge extracellular matrix. Red circles refer to ROIs selected for isotope enrichment observed in a particular image cycle as exemplified by (A) and (B). White circles refer to ROIs, in which an isotope enrichment was observed above or below the displayed image layer. Green circles refer to regions exhibiting no recognizable isotope enrichment within any of the acquired image cycles. This representative image depicts the incorporation of taurine-derived $^{15}N$ and $^{13}C$ into microbial-sized cells as detected by NanoSIMS analysis of extruded extracellular matrix from *I. basta* holobiont samples after incubation with $^{15}N/^{13}C$ labeled taurine over a period of 48 h. Isotope label contents are displayed as isotope fractions on a rainbow color scale, ranging from 0.2 to 1.0 at% $^{15}N$ and 0.9 to 1.6 at% $^{13}C$. The isotope label content of these cells ranged from 0.42–1.15 at% 15N and 1.09–1.47 at% 13C (Fig. S6), which significantly exceeded the values determined within regions of the sponge extracellular matrix (Mann–Whitney U-test, $p < 0.001$), consistent with (a) population(s) of microorganisms specialized in using taurine. Unfortunately, the sponge samples (three explants from the same sponge individual, of which two were used for NanoSIMS analyses) from the taurine incubation experiments could not be analyzed by combining fluorescence in situ hybridization (FISH) with symbiont specific probes and NanoSIMS as this sponge individual exhibited very high autofluorescence at each wavelength typically exploited for the detection of FISH-probe derived signals.

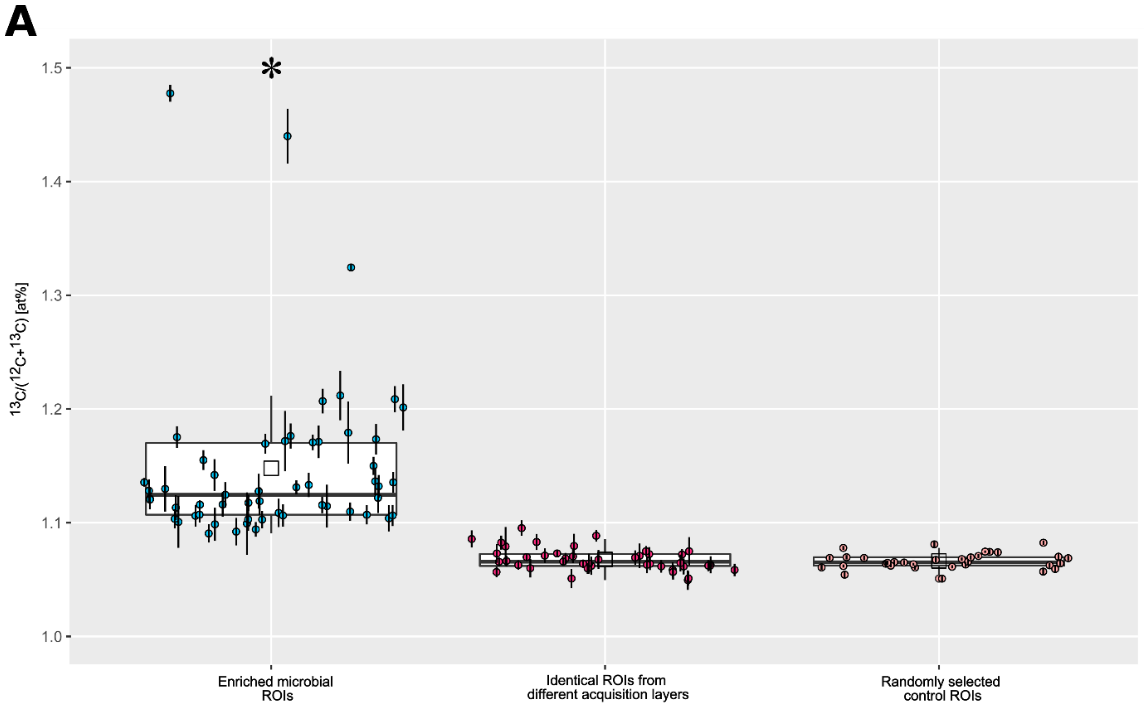


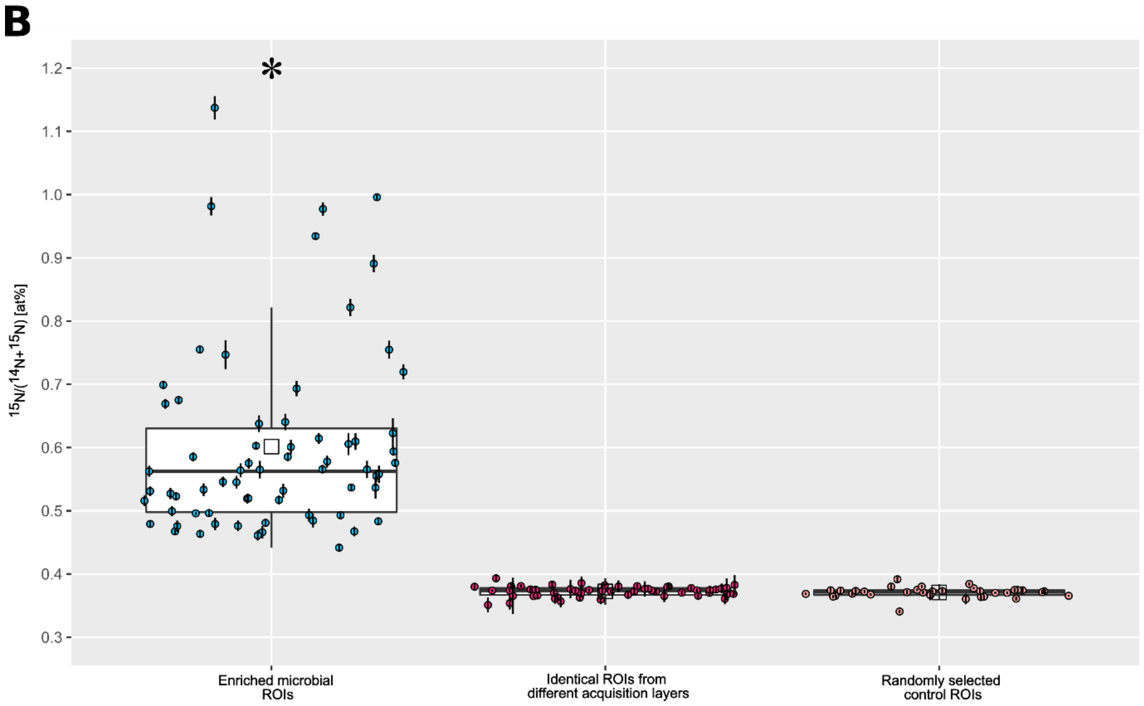


**Figure S6**. Results from region of interest (ROI) specific evaluation of NanoSIMS measurement data from *I. basta* extruded extracellular material. Data acquisition was conducted as multilayer imaging, yielding depth profiles through continuous sputter-erosion by the $Cs^+$ primary ion beam. Each data point refers to the local isotope label content of $^{13}C$ (A) and $^{15}N$ (B) detected within microbial-sized regions. The values displayed on the left refer to ROIs selected for isotope enrichment observed in individual image cycles. The values in the center refer to the values determined in the identical ROIs above or below the image layers indicating isotope enrichment. The values shown on the right correspond to randomly selected microbial-sized ROIs exhibiting no recognizable isotope enrichment within any of the acquired image cycles. The error bars at the individual data points refer to the estimated analytical uncertainty (1σ, Poisson) due to counting statistics (see section Materials and Methods). The definition and classification of ROIs is illustrated in Fig. S5. Boxplots illustrate summary statistics, indicating the median (horizontal bars), first and third quartile (vertical bars: Q1/Q3 + 1.5 × IQR) as well as the mean (open square). Significant differences in isotope enrichment (one-way ANOVA followed by Tukey's test, both $p < 0.001$) are indicated with an asterisk (*).

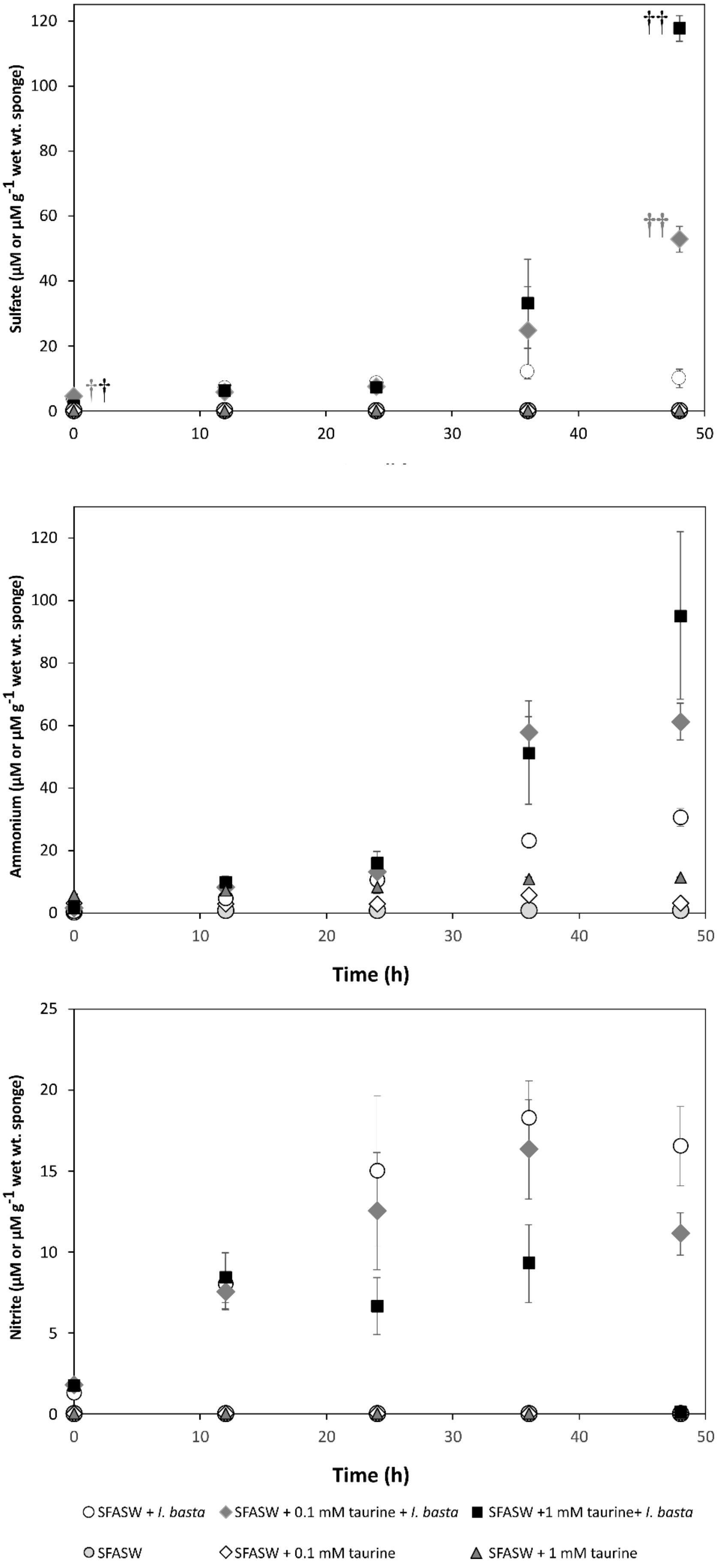

Sulfate (µM or µM g-1 wet wt. sponge)
Ammonium (µM or µM g-1 wet wt. sponge)
Time (h)
Nitrite (µM or µM g-1 wet wt. sponge)
Time (h)
SFASW + I. basta
SFASW + 0.1 mM taurine + I. basta
SFASW +1 mM taurine+ I. basta
SFASW
SFASW + 0.1 mM taurine
SFASW + 1 mM taurine

**Figure S7.** Time series of the concentrations of dissolved sulfate, ammonium, and nitrite in the incubation media of *I. basta* holobiont batch incubations (48 h) in sulfate-free artificial seawater (SFASW) (sampling times at t = 0, 12, 24, 36 and 48 h) performed either with added unlabeled taurine (filled black square: t = 0, +1 mM taurine; filled grey diamond: t = 0, +0.1 mM taurine) or without added taurine (white circle). This resulted in six distinct incubation treatments performed in biological triplicates. The displayed concentrations refer to the measurement values normalized to the wet weight of the respective *I. basta* explants (2.98 ± 0.83 SD g sponge wet wt; range: 1.83–5.28 g sponge wet wt) used in the incubations. The data from the corresponding control incubations with SFASW, but without *I. basta* explants are shown as grey circles for SFASW-only treatments, while SFASW with 0.1 mM taurine and 1 mM taurine treatments are represented by white diamonds and grey triangles respectively. The data for these control incubations are plotted on the same axes as µM. †† symbols in the sulfate concentration panel indicate significant differences for each treatment containing *I. basta* and taurine at t = 48 h (††) relative to at t = 0 (†) (one-way ANOVA followed by Tukey pairwise multiple comparison tests, $p < 0.05$). Nitrite concentrations were found below the limit of detection in all of the incubations conducted in the absence of the *I. basta* holobiont. Error bars refer to +/- 1SE.

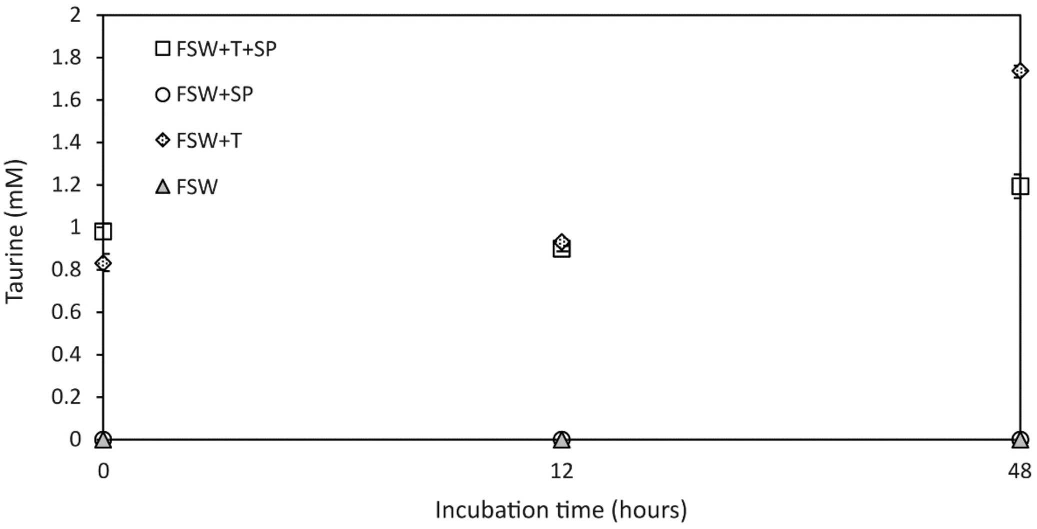


**Figure S8.** Changes in taurine concentrations in seawater after incubation of *Ianthella basta* with 1.6 mM taurine. FSW = natural filtered seawater, FSW+SP = natural filtered seawater conditioned with sponge, FSW+T = natural filtered seawater incubated with taurine (1 mM added at 0 h and 0.6 mM at 36 h) and FSW+T+SP = natural filtered seawater with sponge incubated with taurine (1 mM added at 0 h and 0.6 mM at 36 h). Kruskal-Wallis was used to establish significance. The lower concentration of taurine in FSW+T+SP at 48 h as compared to FSW+T confirms taurine uptake by *I. basta* ($p < 0.05$).

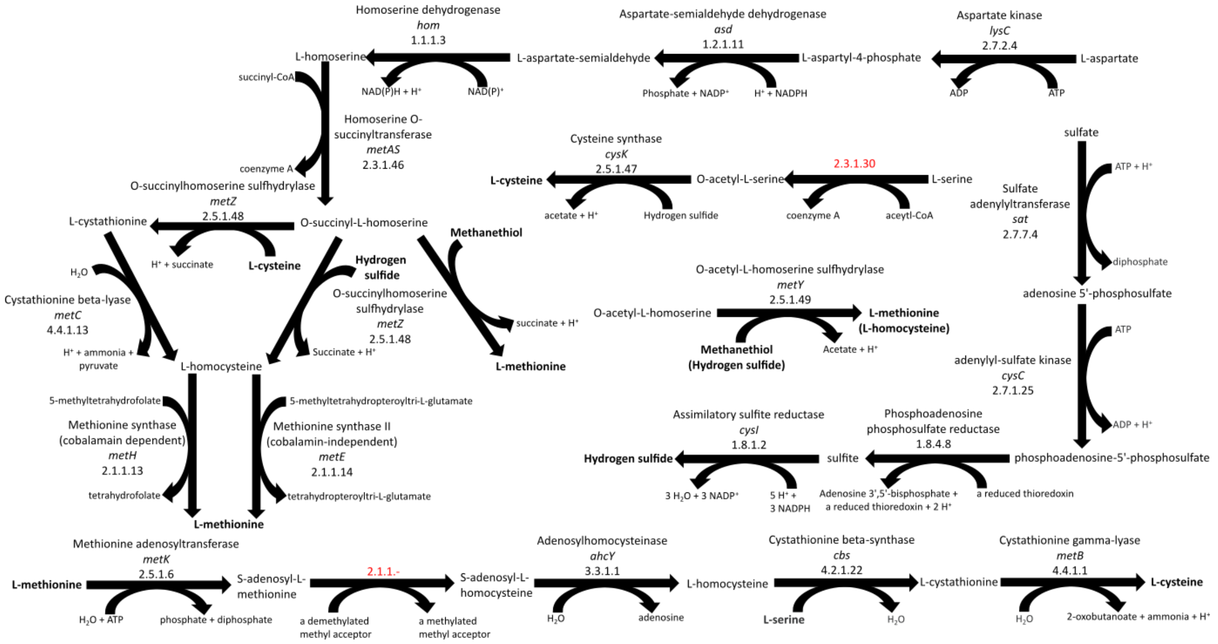


**Figure S9.** Schematic depictions of pathways identified in '*Ca.* Taurinisymbion ianthellae' involved in homocysteine, methionine, and cysteine synthesis. Homoserine synthesis is followed by homocysteine synthesis and culminating in methionine synthesis via either a cobalamin dependent (*metH*), or a cobalamin independent (*metE*) pathway. Alternatively, methionine could be synthesized via methanethiol if supplied as part of the DMSP demethylation pathway as described in Figure 3, which is also cobalamin independent. O-succinylhomoserine sulfhydrylase is a key enzyme (MetZ) that can convert different starting agents either via producing cystathionine when supplied with cysteine, homocysteine when supplied with hydrogen sulfide (from sulfate assimilation — also depicted), or methionine when supplied with methanethiol [12]v(https://enzyme.expasy.org/EC/2.5.1.48). Similarly, MetY, if supplied with O-acetyl-L-homoserine, produces methionine and homocysteine from methanethiol and hydrogen sulfide, respectively [13] (https://enzyme.expasy.org/EC/2.5.1.49). Enzymes labeled in red were not found encoded in the "*Ca.* T. ianthellae" MAG.

**Table S1.** Overview of genome binning parameters, statistics and associated metadata for the IBGammaO2 genome bin of ‘Candidatus Taurisymbion ianthellae’.

| **Bin ID** | **IBGammaO2** |
|---|---|
| Analysis project type | metagenome-assembled genome (MAG) |
| Taxa_id | 16S rRNA and multi-marker phylogenetics |
| Assembly software | Spades (--careful) |
| Annotation | MaGe |
| Genome Quality | High Quality Draft |
| Completeness (%) | 94.82 |
| Contamination (%) | 1.87 |
| Completeness/Contamination Software | CheckM |
| Number of contigs | 97 |
| 16S rRNA gene recovered | yes |
| 16S rRNA gene recovery software | rnammer / MaGe |
| Number of standard tRNAs extracted | 20 |
| tRNA extraction software | tRNA-scan / MaGe |
| Binning software | metabat2 |
| Binning parameters | kmer |
| Genome size (Mbp) | 2.35 |
| N50 (bp) | 41,025 |
| L50 (bp) | 20 |
| Longest contig (bp) | 113,413 |
| Average contig length (bp) | 24,142.92 |
| GC content (%) | 63.49 |
| Protein coding sequences | 2072 |
| rRNAs | 3 |
| tRNAs | 40 |

**Supplementary Note**

Recipes for calcium- and magnesium-free artificial seawater (CMF-ASW) and sulfate-free artificial seawater (SFASW)

| Ingredients | CMF-ASW mM | SFASW mM |
|---|---|---|
| NaCl | 461.85 | 474.33 |
| KCl | 10.73 | 9.00 |
| $CaCl_2$ | - | 9.27 |
| $MgCl_2$ | - | 97.88 |
| $Na_2SO_4$ | 7.04 | - |
| *$NaHCO_3$ | 2.14 | 2.14 |
| $H_2O$ | to 1L | to 1L |

* Add $NaHCO_3$ last.

Adjust to pH 7.8, autoclave, and store at 4°C
